# Impact of preexisting virus-specific maternal antibodies on cytomegalovirus population genetics in a monkey model of congenital transmission

**DOI:** 10.1101/698431

**Authors:** Diana Vera Cruz, Cody S Nelson, Dollnovan Tran, Peter A Barry, Amitinder Kaur, Katia Koelle, Sallie R Permar

**Affiliations:** Computational Biology and Bioinformatics program / Duke Center for Genomic and Computational Biology, Duke University, Durham, North Carolina, USA; Human Vaccine Institute, Duke University School of Medicine, Durham, North Carolina, USA; Tulane National Primate Research Center, Tulane University, Covington, Louisiana, USA; Center for Comparative Medicine, Department of Pathology and Laboratory Medicine, University of California, Davis, California, USA; Department of Biology, Emory University, Atlanta, Georgia, USA

**Author notes:** Address correspondence to Sallie R. Permar.

## Abstract

Human cytomegalovirus (HCMV) infection is the leading non-genetic cause of congenital birth defects worldwide. While several studies have investigated the genetic composition of viral populations in newborns diagnosed with HCMV, little is known regarding mother-to-child viral transmission dynamics and how therapeutic interventions may impact within-host viral populations. Here, we investigate how preexisting CMV-specific antibodies shape the maternal viral population and intrauterine virus transmission. Specifically, we characterize the genetic composition of CMV populations in a monkey model of congenital CMV infection to examine the effects of passively-infused hyperimmune globulin (HIG) on viral population genetics in both maternal and fetal compartments. In this study, 11 seronegative, pregnant monkeys were challenged with rhesus CMV (RhCMV), including a group pretreated with a standard potency HIG preparation (*n* = 3), a group pretreated with a high-neutralizing potency HIG preparation (*n* = 3), and an untreated control group (*n* = 5). Targeted amplicon deep sequencing of RhCMV glycoprotein *B* and *L* genes revealed that one of the three strains present in the viral inoculum (UCD52) dominated maternal and fetal viral populations. We identified *de novo* minor haplotypes of this strain and characterized their dynamics. Many of the identified haplotypes were consistently detected at multiple timepoints within sampled maternal tissues, as well as across tissue compartments, indicating haplotype persistence over time and transmission between maternal compartments. However, haplotype numbers and diversity levels were not appreciably different across HIG pretreatment groups. We found that while the presence of maternal antibodies reduced viral load and congenital infection, it has no apparent impact in the intrahost viral genetic diversity at the investigated loci. Interestingly, some haplotypes present in fetal and maternal-fetal interface tissues were also identified in maternal samples of corresponding dams, providing evidence for a wide RhCMV mother-to-fetus transmission bottleneck even in the presence of preexisting antibodies.

**Author summary:** Human cytomegalovirus (CMV) is the most common infectious cause of birth defects worldwide. Knowledge gaps remain regarding how maternal immunity impacts the genetic composition of CMV populations and the incidence of congenital virus transmission. Addressing these gaps is important to inform vaccine development efforts. Using viral samples collected from a monkey model of congenital CMV infection, we investigated the impact of passively-administered maternal antibodies on the genetic composition of the maternal virus population and that transmitted to the fetus. Our analysis focused on two CMV genes that encode glycoproteins that facilitate viral cellular entry and are known epitope targets of the humoral immune response. By identifying and analyzing variants across sampled maternal tissues, we found no impact in CMV genetic diversity by preexisting CMV-specific antibodies, despite the observation that such antibodies reduce viral load and confer some protection against congenital transmission. We further found that some minor variants identified in fetal and maternal-fetal interface tissues were also present in corresponding maternal tissues, indicating that a large number of viral particles passed from dam to fetus in observed cases of congenital transmission.

## Introduction

Human cytomegalovirus (HCMV) is a member of the β-herpesvirus family and a ubiquitous pathogen that establishes lifelong infection in its host. Seroprevalence rates for HCMV range from 45% in developed nations to 100% in developing nations [1]. While initial HCMV infection is typically asymptomatic in the setting of intact host immunity, congenitally infected infants, immune-compromised individuals, and transplant recipients can suffer adverse HCMV-related outcomes [2–4]. Indeed, HCMV is the leading infectious cause of congenital birth defects, with approximately 1 in 150 live-born infants worldwide infected with HCMV, from which at least 10- 20% will develop long-term sequelae including sensorineural hearing loss, microcephaly, and cognitive impairment [2].

Congenital CMV infection during pregnancy can result from either primary infection or viral reactivation and/or superinfection (secondary infection). While congenital infection could be seeded from the maternal genital tract [5, 6], most cases of transmission are thought to occur from mother to fetus through maternal blood flow to the placenta [7, 8]. High levels of maternal HCMV viremia and maternal infection earlier during gestation have been correlated with a greater risk of congenital infection and more severe congenital disease [9, 10]. Following congenital infection, HCMV can be disseminated throughout the developing fetus with HCMV detectable in multiple fetal tissues in almost 50% of cases [11].

Recent HCMV whole-genome sequencing has revealed that the virus exhibits remarkable genetic diversity both within and between hosts [12–15] despite being a DNA virus. While it is hypothesized that mixed infections and strain recombination are key factors contributing to observed within-host viral diversity [14–16], *de novo* point mutations may also play a role in the generation of intrahost variation. Single nucleotide polymorphisms are distributed unevenly across the viral genome, with more variable regions found within immune evasion-related genes and coding sequences for multiple envelope glycoproteins [15]. Within a single host, these diverse HCMV populations have been observed to change dynamically over time and differ genetically across tissues [12,15,17,18]. Existing studies report the presence of low-frequency intrahost variants following HCMV infection in solid organ transplant recipients and congenital CMV cases [12, 17]. Furthermore, a longitudinal study has shown evidence for persistence of these minor variants over time [12]. And while genetic diversity within a single compartment appears stable over time [13], viral populations from different compartments of a single host can be as genetically-distinct as populations between hosts [13]. Further, viral genomes obtained from the same anatomical compartment across different hosts have been found to show characteristic genetic similarities [15], suggesting that tissue-specific adaptations likely occur and contribute to anatomical compartmentalization.

One of the challenges of HCMV research is that herpesviruses are highly species-specific [19], which has led to a reliance on human clinical trials [7]. Yet, congenital virus transmission can be modeled using both guinea pigs and nonhuman primate models [7, 20]. In particular, rhesus macaques and rhesus CMV (RhCMV) are a highly-relevant model for understanding adult/fetal HCMV pathogenesis [21, 22] and congenital infection [9, 23], as RhCMV is the closest cytomegalovirus species to HCMV [19, 24] and the physiology/immunology of rhesus monkey pregnancy is highly analogous to humans [21]. Previously, our group demonstrated that the depletion of CD4^+^ T cells followed by intravenous RhCMV inoculation of seronegative pregnant monkeys resulted in consistent RhCMV congenital infection and a high rate of fetal loss [24]. We subsequently tested the impact of preexisting antibodies on the incidence and severity of congenital CMV transmission in this monkey model via passive infusion of hyperimmune globulin (HIG) prior to RhCMV inoculation. This study established that preexisting RhCMV-specific antibodies (“standard-potency” HIG) can prevent fetal loss in the absence of functional CD4+ T cell immunity and that highly-neutralizing antibodies (“high-potency” HIG) may block congenital transmission altogether [9]. Furthermore, this previous work demonstrated that potently-neutralizing antibodies present at the time of primary infection can alter viral dynamics *in vivo* [9].

In this study, we focus on the impact of HIG pretreatment on the genetic composition of RhCMV populations found across maternal and fetal tissue compartments. Our analysis is based on RhCMV sequence data derived from maternal compartment samples (plasma, saliva, and urine), samples from the maternal-fetal interface (amniotic fluid and placenta), and fetal tissue samples (fetal heart, brain, lungs, kidney and spleen), where available. Due to the large genome size of RhCMV and a desire to identify viral haplotypes, we focused our approach on amplicon sequencing of variable regions of antibody-targeted glycoprotein genes *gB* and *gL* to explore the effects of preexisting antibodies on viral evolution and tissue compartmentalization. Our work contributes to the deeper understanding of maternal and congenital infection dynamics to better inform developing therapeutic interventions to prevent congenital CMV transmission.

## Methods

### Study setting

Eleven pregnant RhCMV-seronegative dam monkeys were intravenously inoculated with RhCMV to investigate the ability of preexisting antibodies to inhibit congenital CMV transmission. The study consisted of three groups of monkeys: a control group that received no hyperimmune globulin (HIG) pretreatment and two HIG pretreatment groups that differed in their HIG regimen. The first (“standard”) HIG pretreatment group consisted of 3 dams, each of which received a single dose of a standard HIG preparation given 1 hour prior to viral inoculation. The second (“high-potency”) group consisted of 3 dams, each of which received an initial dose 1 hour prior to viral inoculation and a second dose 3 days later. Both doses in the high-potency group used a high-potency HIG preparation by screening serum donor monkeys for serum RhCMV neutralizing activity, as described in [9]. The control group consisted of 5 dams, 3 of which were historical controls [23]. The RhCMV inoculum was a mixture of three different strains: UCD52, UCD59, and 180.92, at relative frequencies of 25%, 25%, and 50% (by infectious viral titer), respectively. These strains are known to have different tropism *in vitro*, with 180.92 isolated on rhesus fibroblasts [25, 26], and UCD52/UCD59 isolated on epithelial cells [27].

For each of the 11 dams studied, samples were taken from maternal blood plasma, urine, saliva, and amniotic fluid at multiple time points following infection. Sample availability varied across dams for reasons such as early fetal loss or low sample volume, previously described in [9]. A subset of the available samples had virus populations that were successfully sequenced and form the basis of our analysis (Table S1). The remainder of these samples did not have successful viral sequencing due to either low viral loads or inadequate sample quality prior to library construction. In addition, virus populations in placental tissue samples from one control group monkey and two standard pretreatment group monkeys, as well as tissue samples from one congenitally infected fetus (from a standard pretreatment group dam) were successfully sequenced (Table S2).

RhCMV viral load was quantified from each sample using qPCR, as described in [9]. For all samples, multiple viral load measurements (3 to 18) were taken to ensure that samples with relatively low levels of virus present were identified as being positive for RhCMV. Viral load on the log10 scale was calculated as the mean of the individual log10 viral load sample measurements. When viral load was below the limit of detection (100 viral copies per ml for plasma and amniotic fluid and 100 viral copies per total DNA µg for urine and saliva), we set its value to half of the detection limit.

### Animal study ethics statement

The animal protocol titled “Maternal immune correlates with protection against congenital cytomegalovirus transmission in rhesus monkeys” was approved by the Tulane University and the Duke University Medical Center Institutional Animal Care and Use Committees (IACUC) under the protocol numbers P0285 and A186-15-06, respectively. Indian-origin rhesus macaques were housed at the Tulane National Primate Research Center and maintained in accordance with institutional and federal guidelines for the care and use of laboratory animals, specifically the USDA Animal Welfare regulations, PHS Policy on Humane Care and Use of Laboratory Animals[28], the NIH/NRC Guide for the Care and Use of Laboratory Animals, Association for Assessment and Accreditation of Laboratory Animal Care accreditation guidelines, as well as Tulane University and Duke University IACUC care and use policies. Tulane National Primate Research Center has strict policies to minimize pain and distress. The monkeys were observed on a daily basis and were administered tiletamine/zolazepam (Telazol), or ketamine if they showed signs of discomfort, pain or distress. In case of illness, the protocol involved analgesics administration and supplemental nutritional support and/or fluid therapy as needed.

Housing conditions were determined by the time and type of RhCMV inoculation, aiming to avoid horizontal transmission of RhCMV from other colony members, where RhCMV is endemic. RhCMV-seronegative pregnant macaques were housed in pairs after RhCMV inoculation if inoculated concurrently with the same viral isolate. Otherwise, single housing in BL2 containment facilities was required. The monkeys were maintained in a standard environment enrichment setting which included manipulable items, swings, food supplements (fruit, vegetables, treats), task-oriented feeding methods as well as human interaction with caretakers and research staff. Dams were released into the colony after 2 or 3 weeks following C-section.

Anesthesia was considered for all procedures considered to cause more than slight pain in humans, including routine sample collection. The agents used included: ketamine, butorphanol, Telazol, buprenorphine, carprofen, meloxicam, and midazolam as needed. The criteria for end-point was defined as loss of 25% of body weight from maximum body weight during protocol, major organ failure or medical conditions unresponsive to treatment and surgical complications unresponsive to immediate intervention. Policies stated that animals deemed at endpoint would be euthanized by overdose of pentobarbital under the direction of the attending veterinarian, consistent with the recommendations of the American Veterinary Medical Association guidelines on euthanasia.

### PCR amplification, viral sequencing. and analysis pipeline

We PCR-amplified two variable regions within the genes encoding RhCMV glycoprotein *B* (gB) and glycoprotein *L (gL)* of RhCMV for next-generation sequencing. The *gB* amplicon was 408 nucleotides long and *gL* amplicon 399 nucleotides long, primer sequences for each amplicon can be found in [9]. Since the RhCMV inoculum consisted of three different strains (UCD52, UCD59, and 180.92), we previously confirmed the absence of primer bias against these strains [9] For each of a given sample’s two amplified loci, our goal was to process two technical replicates. Several samples, however, only had a single successfully sequenced replicate, while other samples had more than two successfully sequenced replicates (Table S1, S2). As described previously [9], each replicate sample was independently PCR-amplified and sequenced following library preparation. Replicates from samples with low viral load were amplified using a nested PCR approach. All replicates were sequenced on an Illumina MiSeq platform, using paired end reads of 300 bases.

To identify viral haplotypes and quantify their frequencies, we first used PEAR [29] to reconstruct (for each available technical replicate) the targeted locus by merging the paired-end reads corresponding to each sequenced fragment. The fused reads were then filtered using the *extractor* tool from the SeekDeep pipeline [30], which filters sequences according to their length, overall quality scores, and presence of primer sequences. Haplotype reconstruction for a given technical replicate was performed on the filtered sequences using the *qluster* tool from SeekDeep, which performs an iterative process of removing spurious, low abundance sequence groups by adding them to more abundant, genetically similar sequence groups when the genetic mismatch between groups occurs at nucleotide positions with low quality.

To obtain a set of haplotypes and their frequencies for a given sample, we combined identified haplotypes across technical replicates. Specifically, for a haplotype to be considered present in a sample, we required it to be detected in both sample replicates. Haplotypes that did not meet this criterion were merged with their genetically-closest haplotype in the sample, and the count of this genetically closest haplotype in the sample was increased accordingly. When only a single replicate was available, we could not perform this step and therefore kept all identified haplotypes present in the single available replicate. When more than two technical replicates were available, we restricted our analyses to the two replicates that were the most similar to one another genetically, based on correlation of haplotype frequencies (see below).

### Quality assurance and error reduction in sequencing data

We performed additional tests and required additional criteria to be met to ensure the quality of each sample that would undergo subsequent analysis. First, to reduce the number of spurious haplotypes in a given sample, we set a frequency threshold that sample haplotypes were required to exceed. This threshold was established as 0.436% based on analysis of plasmid controls. Specifically, we constructed two synthetic plasmids, one containing the *gB* gene and the other containing the *gL* gene. Two technical replicates from each plasmid were sequenced using the same protocol as for the RhCMV samples. Because a single haplotype should be present in these plasmid control populations, any shared low-frequency haplotype is likely a product of PCR amplification error or sequencing error. We found 19 minor haplotypes in the *gB* plasmid control sample after merging technical replicates. These haplotypes ranged in frequency from 0.01% to 0.59% (Figure S1). We found 29 minor haplotypes in the *gL* plasmid control sample after merging technical replicates. These haplotypes ranged in frequency from 0.03% to 0.42% (Figure S1). Our chosen frequency threshold of 0.436% was set at the 0.95 quantile of the combined minor haplotype distributions from the *gB* and *gL* plasmids.

As a second quality assurance step, we performed chimera detection on the haplotypes in each merged sample. A haplotype was classified as a chimera if there was a combination of partial alignments to two observed (and higher frequency) haplotypes in the same sample. Detected chimeras were discarded. These chimeras contributed to only a small fraction of the total reads in each sample (ranging from 1.95% to 7.82% of the reads across all samples).

As a third quality assurance step, we restricted our analysis to those samples that had a Pearson correlation score exceeding 0.70 between the frequencies of the shared haplotypes across technical replicates on the log10 scale. For those samples with only a single technical replicate, we could not perform this step and instead included the sample in our analysis only if the read count exceeded 5000.

Table S1 shows the final set of maternal tissue and amniotic fluid samples that were included in our analyses, for both the *gB* and the *gL* loci. Table S2 shows the set of samples from the maternal-fetal interface (other than the amniotic fluid samples) and from fetal samples that were included in our analyses. In addition to these samples, the genetic composition of the inoculum was analyzed. Each of the three viral stocks comprising the inoculum (UCD52, UCD59, 180.92) was independently sequenced. Two successfully sequenced replicates were available for each of the three stock samples.

### Strain classification and nucleotide diversity calculations

Each identified haplotype in a sample was classified as belonging to one of the three strains that comprised the inoculum (UCD52, UCD59, or 180.92) based on its genetic distance to the reference sequences of these three strains. The reference sequences of the targeted *gB* and *gL* regions were obtained from [27] for strains UCD52 and UCD59 and from [26]. for strain 180.92. Nucleotide diversity *π* present in a sample was calculated for each strain independently using the commonly used Nei-Gojobori equation, as described in [31]. All identified haplotypes, across all sequenced samples, are listed in the **Appendix S3 and Appendix S4**. The frequencies of these haplotypes in each of the sequenced samples, including the inoculum, are given in the **Appendix S1 and Appendix S2**.

### Statistical analysis and software

Data processing, analysis, and visualization were performed in R. Pairwise comparisons between groups were performed using non-parametric tests as indicated. For the network visualization of haplotypes, we employed the R package *RCy3* version 1.2.0 that interfaces R 3.4 with Cytoscape.

### Data and code availability

Sequencing data in fastq format from all the samples is available in SRA under the Bioproject PRJNA386504. Primer sequences for the gB and gL regions can be found in [9]. All the R codes required for this study are available on GitHub: dverac/SNAPP.

## Results

### Maternal viral load dynamics, congenital transmission, and strain dominance

As previously described [9], dams in the high-potency HIG pretreatment group had reduced peak viral loads in maternal plasma relative to dams in the control group following primary maternal infection (Figure S2A). Viral kinetics in the saliva and urine were also delayed in the high-potency pretreatment group compared to the control group (Figure S2C,D) [9]. Interestingly, copies/mL transmitted the virus to the amniotic fluid compartment. This included all 5 dams in the control group, 2 out of 3 dams in the standard pretreatment group, but none of the 3 dams in the high-potency pretreatment group. Viral dynamics in the amniotic fluid, when present, did not appear to differ between the control group monkeys and the standard HIG pretreatment group monkeys (Figure S2B).

Of the three viral strains used in the RhCMV inoculum, UCD52 became dominant in the overwhelming majority of tissue compartments, regardless of pretreatment group status (Figure S3) [9]. The single exception to this, which was supported by both the *gB* and *gL* loci, was a week 1 plasma sample from a dam from the control group (C1) in which UCD59 haplotypes were the most abundant. Given the dominance of the UCD52 strain in the overwhelming majority of samples, we focused our remaining analyses on haplotypes that were classified as belonging to the dominant UC52 strain.

### Minor RhCMV haplotypes and levels of genetic diversity during acute maternal infection

Across the majority of analyzed samples, we found that the dominant *in vivo* UCD52 haplotype was the canonical UCD52 reference haplotype of the viral inoculum. This was the case both for the *gB* locus and the *gL* locus, and across all groups and compartments studied.

Our analysis of amplified sequences from the *gB* locus identified a large number of minor haplotypes in maternal and fetal compartments that differed from the canonical UCD52 *gB* haplotype by typically only a single nucleotide (Figure 1, Figures S4-S11). These minor haplotypes ranged in frequency from just above the sequencing error cut-off frequency of 0.436% up to 43.27%, with a median frequency of 0.80%. Maternal samples differed in the number of identified *gB* haplotypes they contained, ranging from 1 to 33, with a median of 5 haplotypes per sample. The number of haplotypes identified in a sample was not positively correlated with the sample’s viral load (Figure S12), indicating that the numbers of observed haplotypes were not restricted by sample viral load. Given our constrained cut-off for haplotypes detection, these minor haplotypes are potentially produced *de novo* as RhCMV spreads within each monkey. Within individual dams, we observed that some of the minor haplotypes were shared across timepoints from the same compartment and/or across compartments (Figure 1, Figures S4-S11). This finding indicates that some of these minor haplotypes persist over a timespan of weeks in a given compartment and that some of these minor haplotypes are likely transmitted across anatomic compartments. Of the minor haplotypes that were shared across compartments, most were shared between the plasma and one other compartment (Figure 1, Figure 2, Figures S4-S11). This pattern may reflect plasma being a source of viral haplotypes for other compartments; alternatively, it may simply be due to a larger number of plasma samples being successfully sequenced relative to those from other compartments (Table S1). Interestingly, in 6 out of the 8 monkeys that had both urine and saliva sequences available, there were also minor *gB* haplotypes that appeared to be shared exclusively between urine and saliva samples. These haplotypes were generally found first in urine and then in a later week in the saliva, suggesting potential oral auto-inoculation from virus shed in urine.

**Figure 1.**
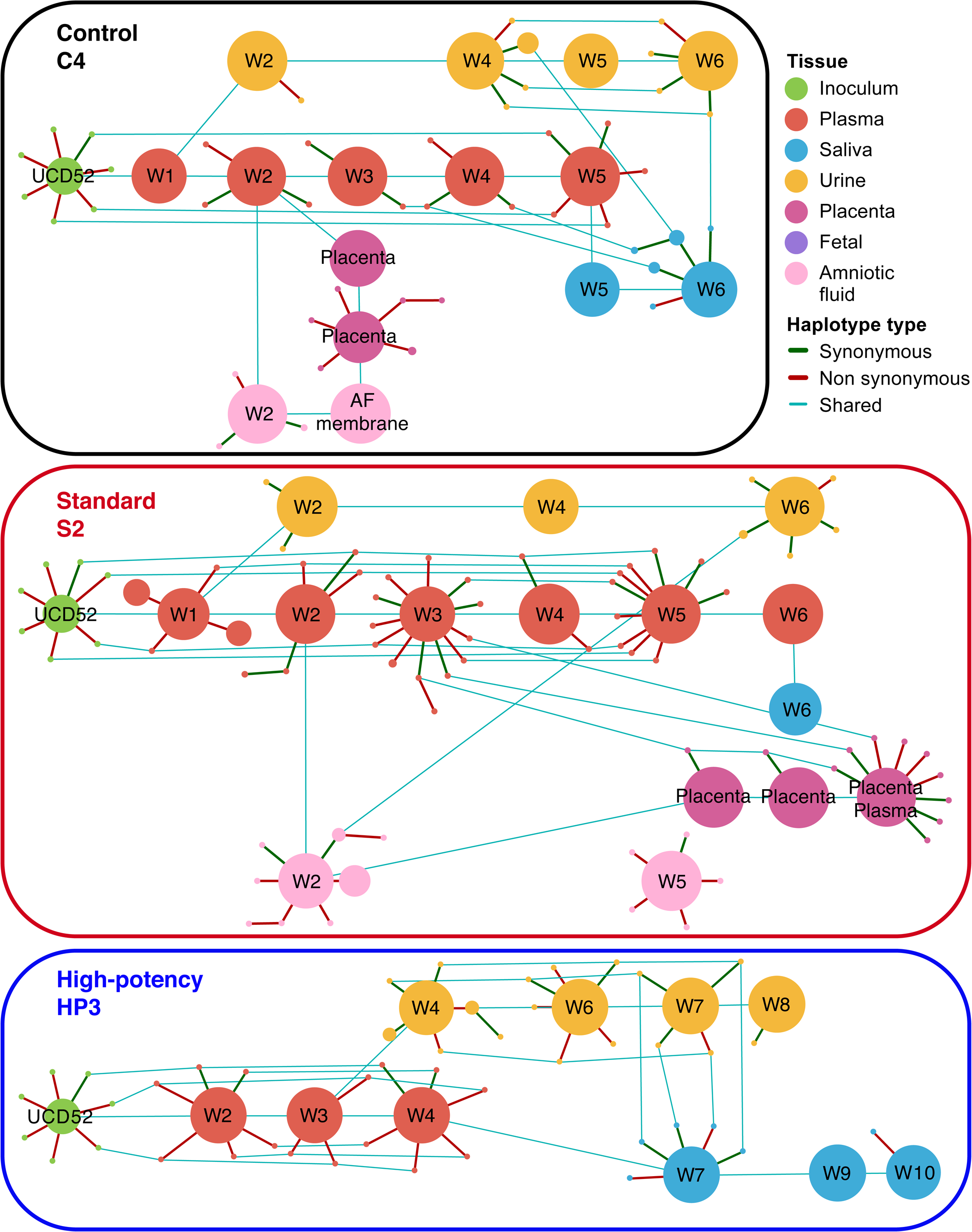
UCD52 haplotype networks for the *gB* locus across sampled tissues from three representative monkeys. Haplotype networks are shown for one control group dam (A), one standard pretreatment group dam (B), and one high-potency pretreatment group dam (C). Edges connect haplotypes that differ by a single nucleotide, with green edges depicting synonymous mutations and red edges depicting nonsynonymous mutations. Node sizes scale with haplotype relative frequency. Samples are labeled by collection week. Blue lines connect shared haplotypes across samples.

**Figure 2.**
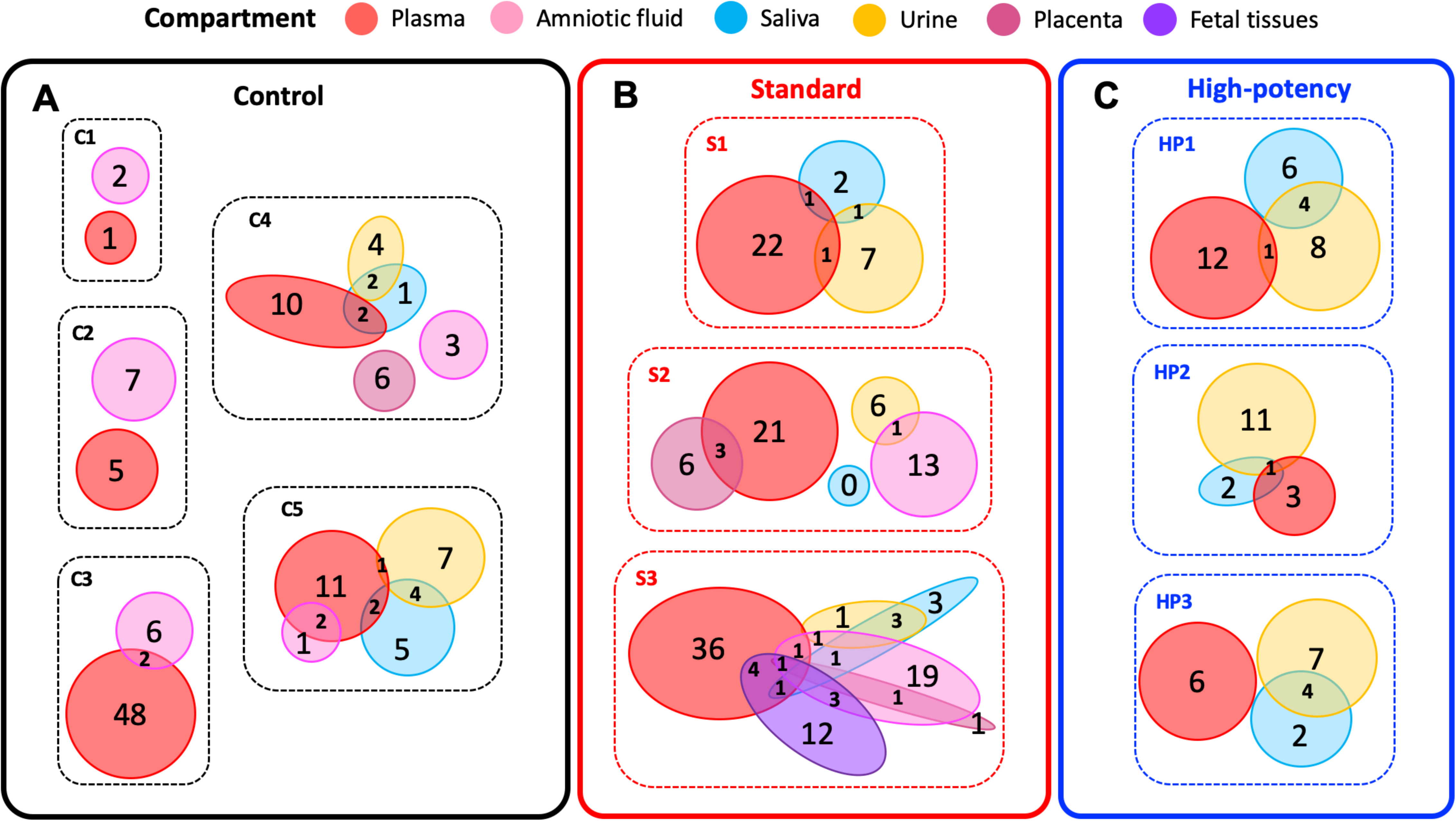
The number of UCD52 minor *gB* haplotypes that are either shared or unique across compartments, by monkey. Here, the set of minor haplotypes for a given compartment includes all timepoint samples from that compartment. Patterns of minor haplotype sharing for (A) control group monkeys, (B) standard pretreatment group monkeys, and (C) high-potency pretreatment group monkeys. Compartments are color-coded as in Figure 1.

To assess whether the number of identified *gB* haplotypes differed by pretreatment group, we calculated the median number of minor UCD52 haplotypes in each available tissue for each of the 11 dams. We found no significant differences in the median number of minor *gB* haplotypes by tissue across any pair of pretreatment groups (all Mann-Whitney *U* tests > 0.1; Figure 3A). To determine whether certain tissues tended to allow more non-synonymous variation than other tissues, or whether the extent of nonsynonymous variation differed by pretreatment group, we further calculated the proportion of minor *gB* haplotypes that differed from the canonical haplotype by a nonsynonymous mutation, by tissue and monkey. No major differences were found between tissues or between pretreatment groups (for data on haplotypes, see **Appendix S1**).

**Figure 3.**
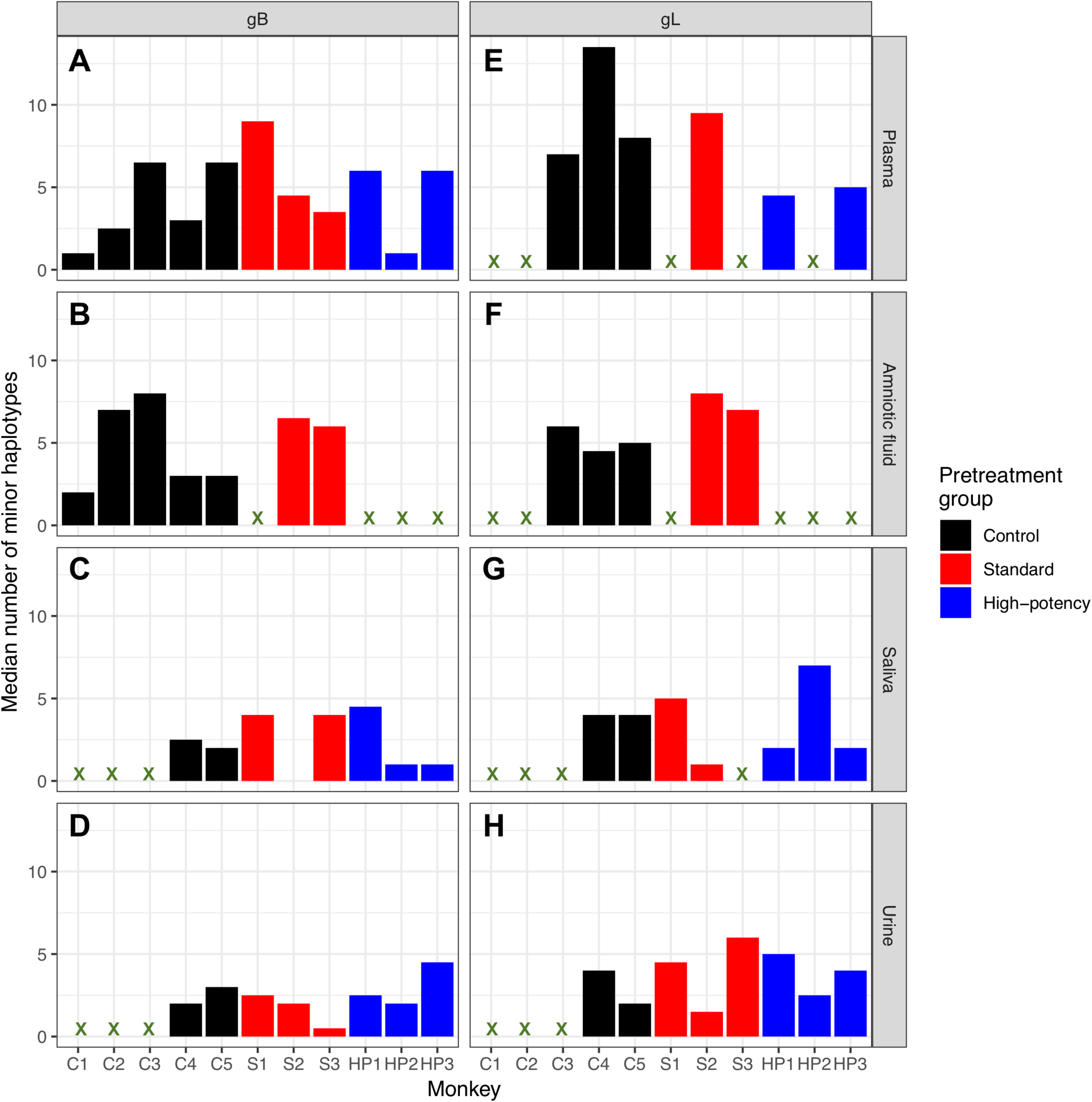
Median number of minor haplotypes, by locus, tissue, and pretreatment group. The left column shows the median number of *gB* minor haplotypes; the right column shows the median number of *gL* minor haplotypes. Rows show tissues: plasma, amniotic fluid, saliva, and urine. Marker symbols correspond with those in Figure S2.

The UCD52 haplotype patterns observed using the *gL* locus are consistent with those at the *gB* locus. Specifically, minor *gL* haplotypes generally differed from one of the two dominant *gL* haplotypes present in the inoculum by a single nucleotide (Figure S13-S22). Similar to the frequencies observed for *gB* haplotypes, minor *gL* haplotypes were present at frequencies as low as 0.44% and up to 48.16%, with a median frequency of 1.05%. Samples differed in the number of identified *gL* haplotypes they contained, ranging from 2 to 29 with a median of 6 haplotypes per sample. Again, no correlation was found between the number of haplotypes identified in a sample and the sample’s viral load (Figure S23). Some of the identified minor *gL* haplotypes appeared to persist within the same tissue over time, and some were shared across tissue compartments. Similar to our findings at the *gB* locus, most of the minor haplotypes that were shared across compartments were shared between the plasma and one other compartment (Figure S24). Minor *gL* haplotypes shared between urine and saliva compartments again suggested auto-inoculation. Finally, consistent with the findings from the *gB* locus, the median number of *gL* minor haplotypes observed in any tissue did not differ between pretreatment groups (Figure 3). We again found no significant differences between tissues or pretreatment groups in the proportion of minor *gL* haplotypes that were nonsynonymous (for data on haplotypes, see **Appendix S2**), consistent with the lack of pattern at the *gB* locus.

We next assessed whether HIG pretreatment had an impact on RhCMV genetic diversity, as measured by pairwise nucleotide diversity *π* for each sample’s UCD52 viral population. Levels of viral genetic diversity varied significantly between monkeys, compartments, and across weeks (Figure 4 for *gB*, Figure S25 for *gL*). Despite this variation, median levels of *gB* viral genetic diversity did not differ by pretreatment group for any tissue (all Mann-Whitney U tests > 0.1) besides the amniotic fluid. In this compartment, median levels of *gB* viral genetic diversity appeared to be slightly higher in standard pretreatment group monkeys than in control animals (Mann-Whitney U test, *p* = 0.05). Median levels of *gL* viral genetic diversity did not differ by pretreatment group for any tissue (all Mann-Whitney U tests > 0.1; Figure S25).

**Figure 4.**
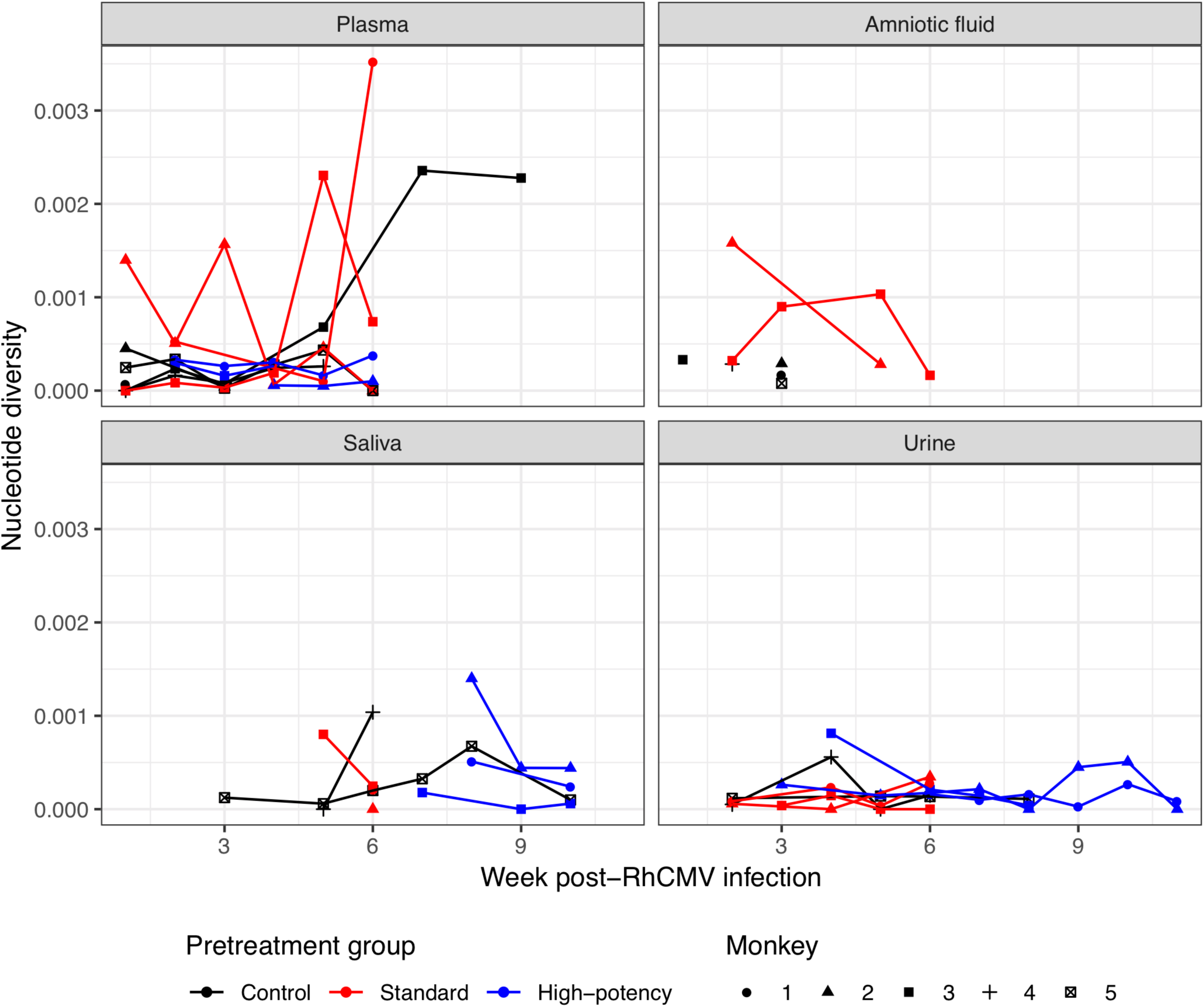
Pairwise genetic diversity *π* over time, by tissue, for the *gB* locus. Marker symbols correspond with those in Figure S2.

### Genetic diversity and compartmentalization of maternal RhCMV variants identified in placenta and amniotic fluid

We next sought to determine the extent to which minor UCD52 haplotypes were shared between maternal compartments and compartments comprising the maternal-fetal interface (amniotic fluid and placental tissues). As reported above, we found that some minor *gB* and *gL* UCD52 haplotypes were shared between maternal plasma samples and amniotic fluid samples (Figures 1 and 2; **Appendix S1** and **Appendix S2**). Similarly, some of the minor *gB* and *gL* UCD52 haplotypes found in placental tissue samples were also present in maternal plasma samples (Figure 2). Specifically, between placental tissues and plasma samples, we observed 6 shared minor *gL* haplotypes in C4 (Figure S15), 3 shared minor *gB* haplotypes in S2 (Figure 1), 1 shared minor *gL* haplotype in S2 (Figure S18), and 1 shared minor *gB* haplotype in S3 (Figure S9). As these minor shared haplotypes are mostly present at marginal frequencies in maternal tissues (median frequency in plasma for shared haplotypes: 1.28%, minimum 0.47%, in gB, S2; maximum 24.05% in gL, S2) (**Appendix S1** and **Appendix S2**), the bottleneck between mother and fetus is likely relatively large. Interestingly, we observed similar or larger number of minor *gB* and *gL* UCD52 haplotypes in amniotic fluid samples compared to placental tissues (Figure 2, Figure S24; **Appendix S1** and **Appendix S2**), indicating that *de novo* viral mutations may occur in the fetus and subsequently be shed into the amniotic fluid. Interestingly, in the one case in which placental plasma was available for analysis (dam S2 in Figure 1; Figure S18), we found considerably more minor haplotypes in both *gB* and *gL* gene regions in this sample compared with placental tissue samples and many of these minor haplotypes were not observed in maternal plasma samples.

### Genetic diversity and compartmentalization of fetal RhCMV variants

Congenital infection was confirmed in two of three dams in the standard pretreatment group and in all five control dams. Yet, all five control dams experienced fetal loss within 2-3 weeks of maternal infection and fetal tissues were often not recovered. In standard pretreatment group dam S3, nearly all the fetal tissues harvested at 6 weeks post-RhCMV infection tested positive for RhCMV, including fetal lung, brain, kidney, spleen, heart, placenta, amniotic fluid, and amniotic membrane. Similar to our observation in the maternal tissue compartments, a single major UCD52 haplotype (the canonical reference haplotype) was present in all fetal tissue samples. Multiple minor UCD52 haplotypes were also detected in these samples (Figure S9; Figure 5). Intriguingly, a second, minor haplotype was found in each of the fetal tissues, present at frequencies ≤1%. This haplotype was also observed in one of the paired dam’s three amniotic fluid samples (week 3, frequency of 0.8%), placenta (frequency of 0.85%), and in two of the paired dam’s plasma samples (at weeks 3 and 6; frequencies of 0.58% and 0.66%, respectively) (Figure 5). Of the remaining 21 minor UCD52 haplotypes in fetal tissues, 4 were also present in amniotic fluid samples and 5 in plasma samples of paired dam S3 (Figure 5). Plasma haplotypes detected as late as weeks 5 and 6 post-inoculation contribute to those shared haplotypes. In comparison to the 10 minor haplotypes shared between fetal tissues and maternal plasma/amniotic fluid samples from the paired dam S3, only 0-5 minor haplotypes were shared between the fetal tissues and non-paired dams. We then calculated pairwise genetic diversity *π* from each available fetal tissue. We observed lower diversity in the fetal tissues compared to that in both the amniotic fluid (*p* = 0.025) and the plasma at late weeks post-infection (Weeks 4 to 6, *p* = 0.095). We further observed higher diversity in the fetal tissues compared to that in plasma during the first three weeks post-infection (Weeks 1 to 3, *p* = 0.024). These observations together suggest that the maternal viral population contributes to the viral diversity in the fetus and that congenital transmission may be subject to a wide and potentially continuous bottleneck, through which various minor haplotypes present in maternal plasma or amniotic fluid can be transmitted to the fetus.

**Figure 5.**
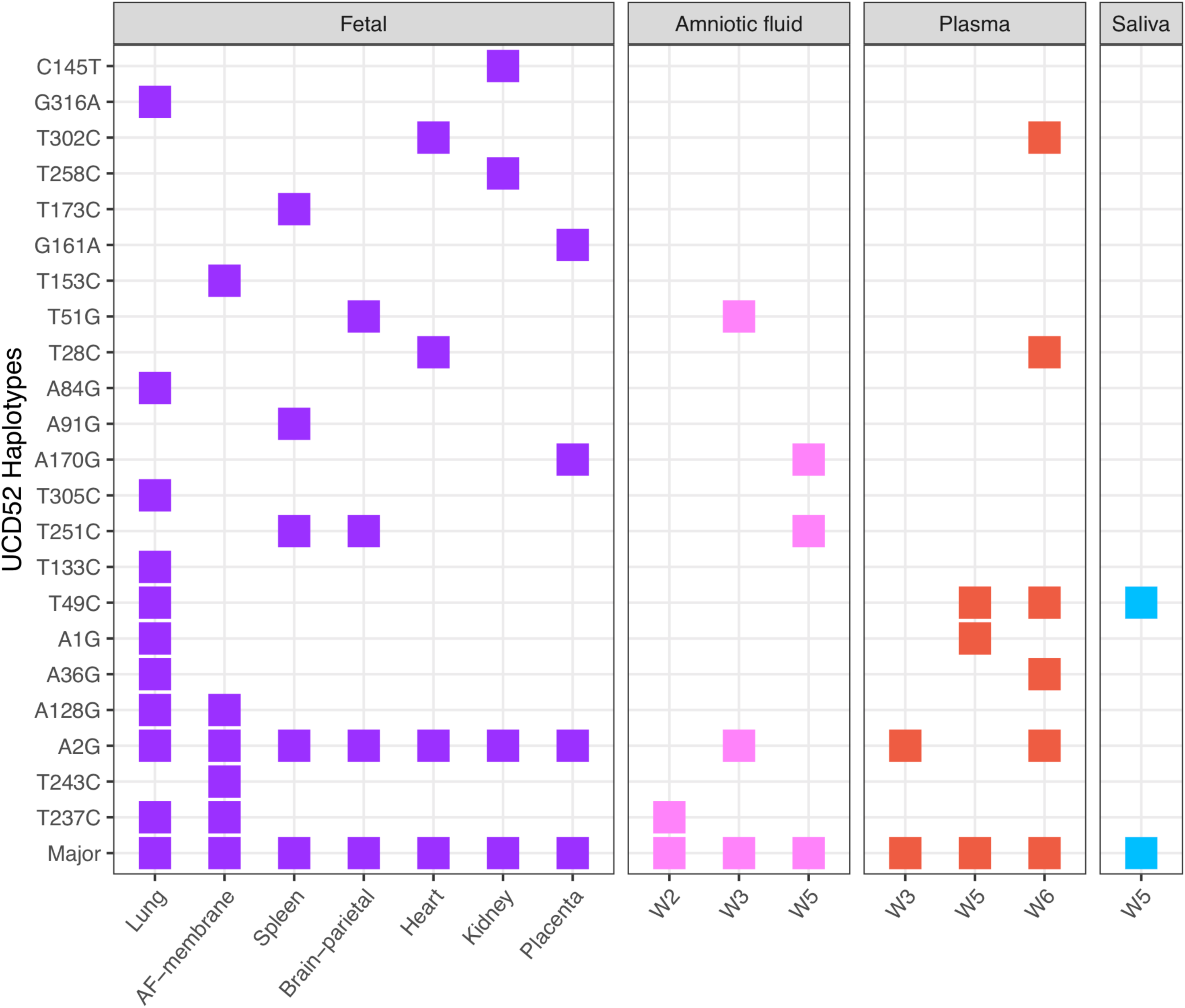
Minor UCD52 haplotypes found in fetal tissues, and their presence in maternal compartments of dam S3. Each row depicts a haplotype found in at least one fetal tissue (purple), harvested at 6 weeks post-RhCMV infection. Rows are ordered from highest frequency haplotype (bottom row) to lowest frequency haplotype (top row).

## Discussion

In this study, we characterized the population genetics of RhCMV in a nonhuman primate model of congenital CMV transmission and quantified the impact of preexisting maternal virus-specific antibodies on the viral population at the maternal-fetal interface. Unique aspects of this study include serial sampling from multiple maternal compartments over the time period of an acute RhCMV infection as well as the sequencing of RhCMV-infected tissues at the maternal-fetal interface and, in one instance, from fetal tissues. We found that all maternal and fetal tissue samples (excepting one) were dominated by UCD52. Furthermore, there was a trend towards higher UCD52 haplotype frequencies in the plasma of high-potency pretreatment group monkeys compared to those in the control group monkeys and standard pretreatment group monkeys. The reason for the dominance of this singular strain across HIG pretreatment groups is unclear, although it likely indicates that the UCD52 strain is more genetically fit for *in vivo* replication than either of the co-inoculated variants UCD59 and 180.92. Given the dominance of UCD52 in all groups and tissues, we focused subsequent analyses on characterizing the genetic variation of this strain in available samples. In the majority of samples from maternal tissues, maternal-fetal interface, and fetal tissues, the major *gB* and *gL* UCD52 variant detected was the canonical UCD52 reference sequence. However, most samples also had low-frequency (minor) UCD52 haplotypes present, with some of these minor haplotypes persisting over time and occasionally shared between sampled compartments.

In our analyses, despite high-potency pretreatment monkeys having significantly lower peak viral loads compared to standard and control group monkeys, we found no strong evidence for a relationship between HIG pretreatment and maternal plasma UCD52 haplotype number or nucleotide diversity. Together, these results suggest that preexisting antibodies can reduce overall viral load but do not appear to restrict replication of specific UCD52 viral variants or limit viral diversity. We also found no significant differences in saliva or urine virus haplotype number or nucleotide diversity between the three groups at either *gB* or *gL* loci. While we previously assessed and reported lower *maximum* plasma viral diversity levels in monkeys pretreated with HIG compared to the control group [9], here, we included a more in-depth analysis across timepoints to report the *median* viral diversity levels across monkeys, and did not find any lasting impact of preexisting antibodies on maternal viral diversity.

Our identification of shared, minor UCD52 haplotypes between maternal plasma samples, amniotic fluid, placental tissue, and fetal tissues is consistent with previous studies investigating the population genetics of HCMV in newborns [12, 32], which together point towards a loose vertical transmission bottleneck between mother and fetus. While previous studies have estimated transmission bottleneck sizes for HCMV and other viruses, in this study we were unable to quantify transmission bottleneck sizes between mother and fetus due to: 1) low levels of haplotype diversity and 2) haplotype frequencies near the limit of detection. Nevertheless, based on the identification of minor UCD52 haplotypes across maternal, maternal-fetal interface, and fetal tissues, our analysis suggests that diversity in a given tissue is generated through a combination of multiple viral haplotypes being passed to that compartment, along with *de novo*, local generation of viral mutations.

Recently, Sackman and coauthors proposed a model for congenital human CMV (HCMV) transmission that involves two successive transmission events: maternal virus infection of placental tissues followed by continued transmission of the placental viral population to fetal circulation [33]. This model is supported by observations of the sustained presence of HCMV in the placenta and umbilical cord, which would potentially allow for transmission between placental tissues and fetal tissues over a longer time interval [34]. Our results provide further support for this proposed model. Specifically, in our analysis of fetal samples from dam S3, we identified minor haplotypes in fetal tissues that were also present in maternal plasma (Figure S9). Furthermore, we observed that multiple haplotypes in this dam’s amniotic fluid were also observed in maternal plasma and other maternal compartments. Since amniotic fluid haplotypes derive from both intrauterine and fetal viral populations, this finding again provides support for a loose transmission bottleneck from mother to fetus.

Our conclusions are limited by multiple factors. First, as is common for experimental monkey challenge studies and particular to studies of a selective colony of RhCMV seronegative breeding animals, we are limited by the small number of animals in each group and by sample availability. PCR amplification failure further limited the number of samples available for analysis (Table S1). Second, this study did not employ full genome sequencing, but instead sequenced only two gene regions (*gB* and *gL*) to allow for studies of virus population in samples with low viral load. Subsequently, any effect of antibody selection over the non-sequenced regions of gB and gL or over other viral proteins will not be observed. Third, because RhCMV is a DNA virus with a low mutation rate, our conclusions were limited by the low levels of genetic diversity observed in the samples. We also used highly conservative haplotype-calling and error reduction methods to ensure that the haplotypes we identified were not false positives. As a result, however, we likely excluded many true haplotypes, which reduced the diversity levels we characterized and limited our ability to make inferences regarding transmission bottleneck sizes. Finally, given that our animal model of congenital CMV transmission involves maternal CD4^+^ T cell depletion, which results in consistent placental transmission, our results might not be applicable to immunocompetent individuals.

Despite these limitations, however, we were able to conclude that minor haplotypes persisted over time within single maternal tissue compartments and that these minor haplotypes were occasionally shared between anatomic compartments. Moreover, there was not a strict bottleneck for the viral major and minor haplotypes that appeared in placenta, amniotic fluid, and fetal tissues. All these observations are consistent with those from human congenital CMV cases [12,15,35–37]. Patterns of viral diversity within and across compartments, however, did not appear to differ between HIG pretreatment groups. These findings indicate that, although potently-neutralizing CMV-specific antibodies can effectively reduce viral population size and prevent congenital transmission [9], preexisting HIG had limited impact on the genetic makeup of the maternal RhCMV populations or transmitted variants. These findings are interesting given the growing evidence that preexisting HCMV-specific antibodies can reduce the incidence and severity of congenital HCMV [9,38–40], perhaps suggesting a model wherein congenital virus transmission is dependent upon the overall quantity of maternal systemically-circulating virus rather than antibody selection of specific variants at the maternal-fetal interface. Further studies, ideally starting with an inoculum containing higher levels of viral diversity, may be required to provide a deeper understanding of the extent of antibody-mediated immune-pressure on CMV populations, as well as the effect of antibodies on viral transmission dynamics across the placenta. Results from these studies will be critical to more effectively anticipate the effect of CMV vaccines and therapeutic interventions on congenital CMV transmission potential and the propensity for this virus to evolutionarily circumvent these interventions.

## Author contributions

A.K., K.K. and S.R.P. designed research; C.S.N., D.T. and D.V.C. performed research; D.V.C. analyzed data; P.A.B. and K.K. contributed analytic tools/expertise/reagents; and D.V.C, C.S.N., K.K. and S.R.P. wrote the paper.

## Acknowledgements

This work was supported by NIH/NICHD Director’s New Innovator grant to S.R.P (DP2HD075699), NIH/NIAID grants to S.R.P. and K.K. (R21AI136556), fellowship grant to C.S.N (F30HD089577), and NIH P51 OD011104 to the Tulane National Primate Research Center. The funders had no role in study design, data collection and interpretation, decision to publish, or the preparation of this manuscript. The content is solely the responsibility of the authors and does not necessarily represent the official views of the National Institutes of Health.

## Supplementary figures legends

**Figure S1.**
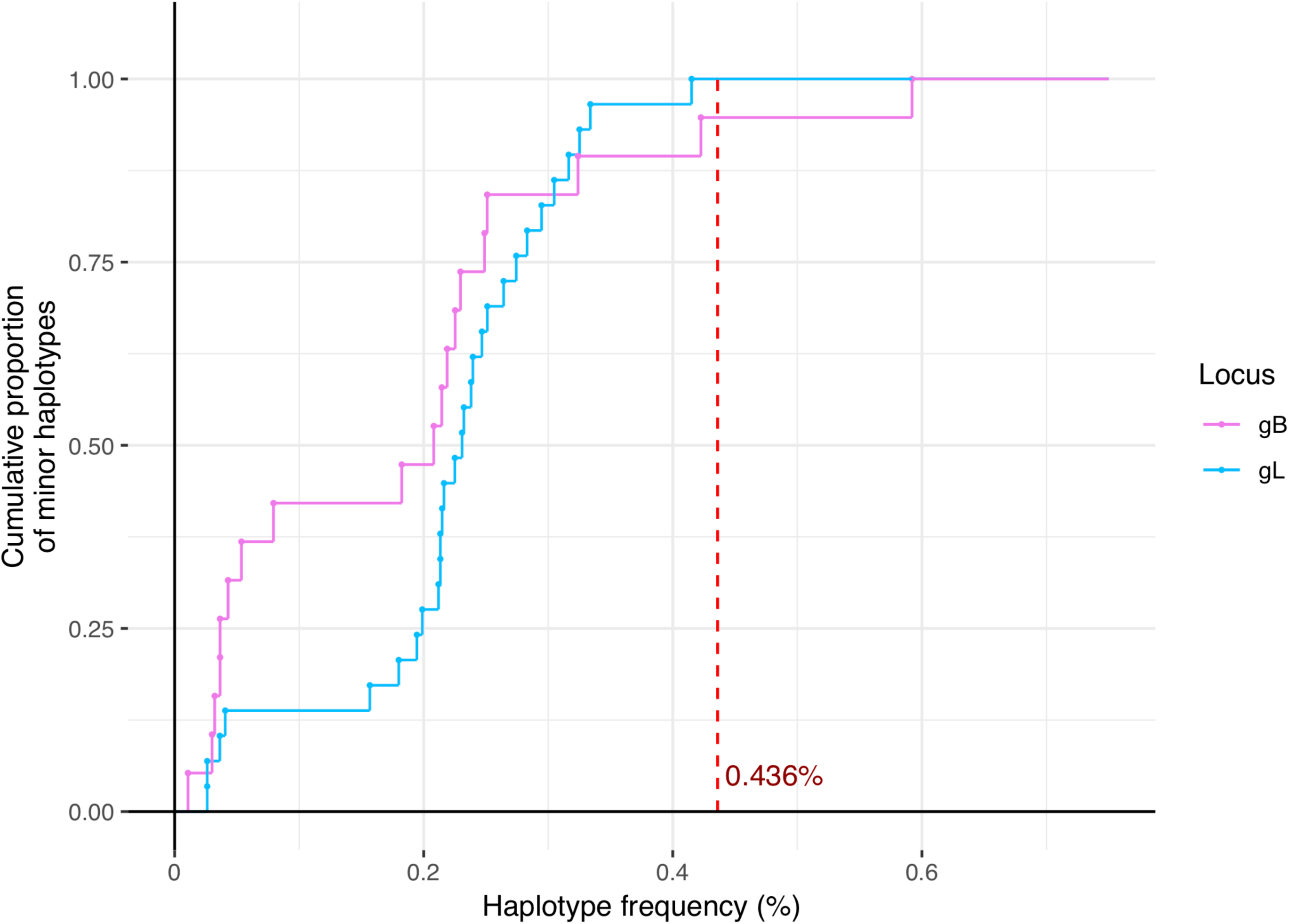
The use of synthetic plasmids to define a frequency threshold to exclude spurious haplotypes from samples. Minor haplotypes were identified from the synthetic plasmid control samples as described in the Methods section, for both the *gB* locus and the *gL* locus. The figure shows, for each locus, the fraction of identified minor haplotypes (y-axis) that fall at the haplotype frequency shown on the x-axis or below. The vertical red line shows the frequency threshold of 0.436% that was used to call minor haplotypes.

**Figure S2.**
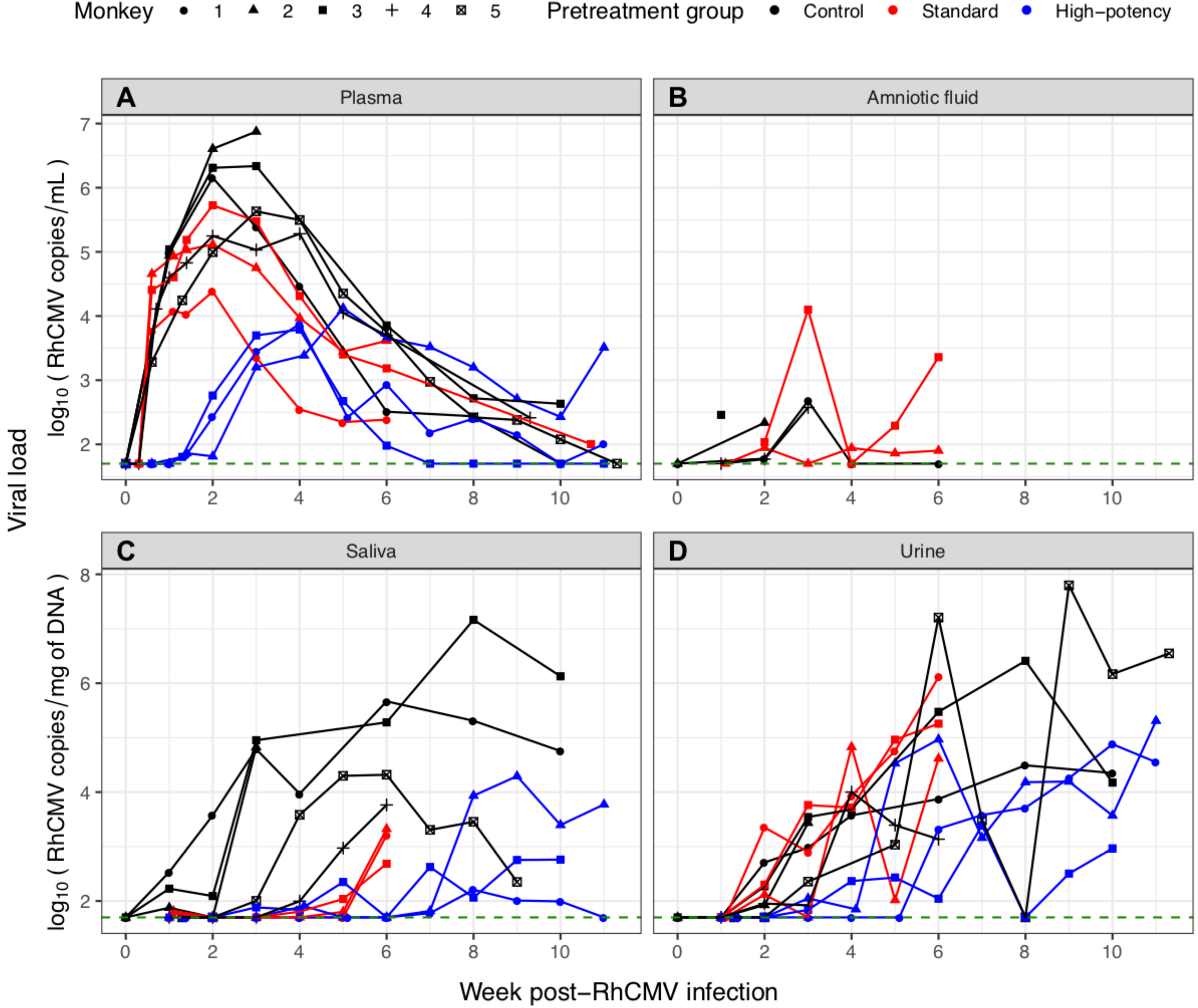
Viral load dynamics measured in dams experimentally infected with RhCMV. Virus was measured in (A) plasma, (B) amniotic fluid, (C) saliva, and (D) urine. Monkeys are color-coded according to pretreatment group: control (black), standard pretreatment group (red), and high-potency pretreatment group (blue). Monkey ID numbers correspond to those provided in Table S1. Virus was detected in the plasma, saliva, and urine of all 11 monkeys. Virus was only detected in the amniotic fluid of the 5 control group monkeys and in 2 of the 3 standard HIG group monkeys. Viral load levels shown here are average values when more than one measurement was available (Methods).

**Figure S3.**
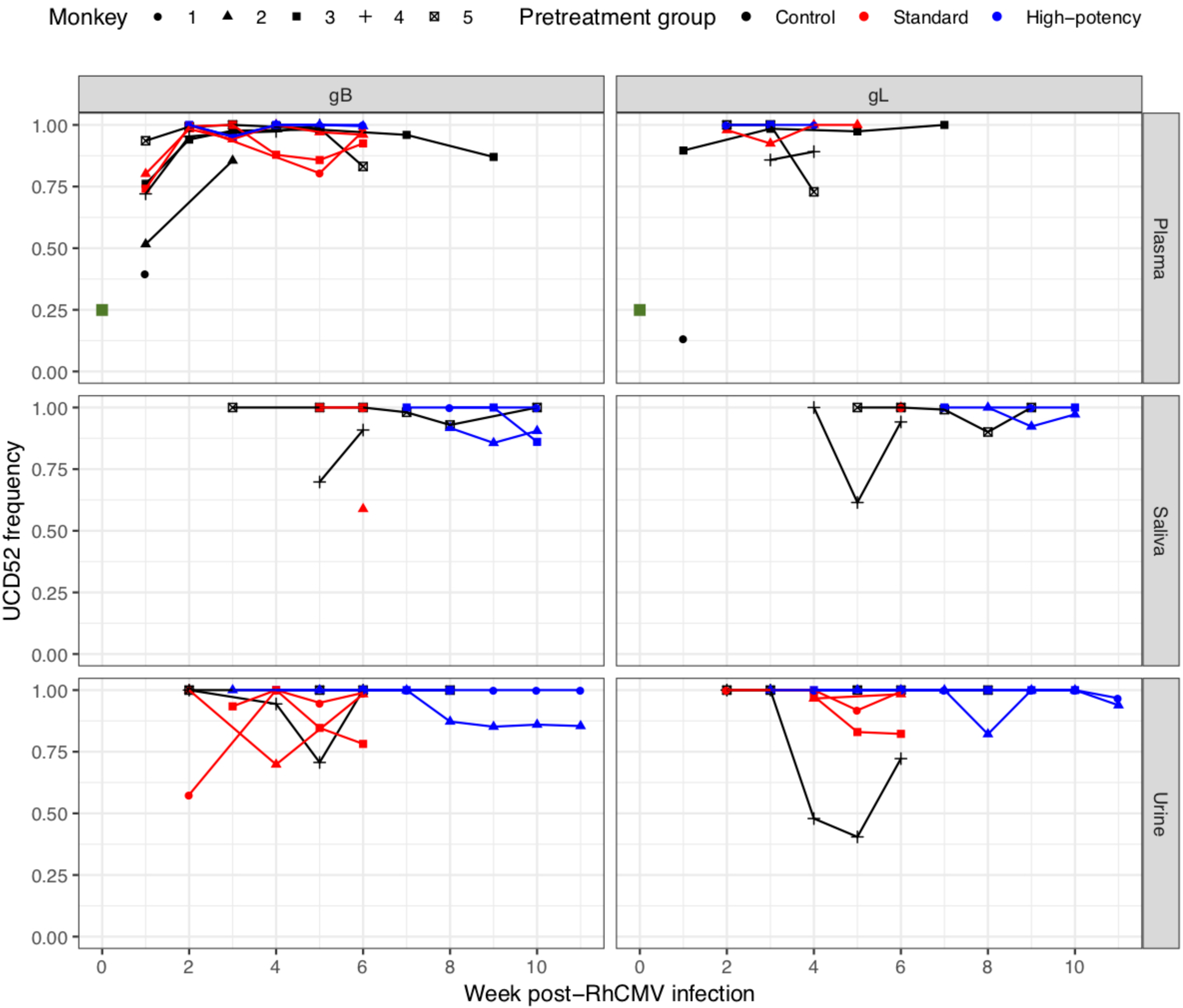
Strain composition of the RhCMV population in various maternal compartments over time. The proportion of the RhCMV population belonging to strain UCD52 is shown for maternal plasma, saliva, and urine. Strain frequencies were calculated for the *gB* locus (left column) and for the *gL* locus (right column). Green squares in the plasma subplots denote the fraction of the viral inoculum that was UCD52 (25%).

**Figure S4.**
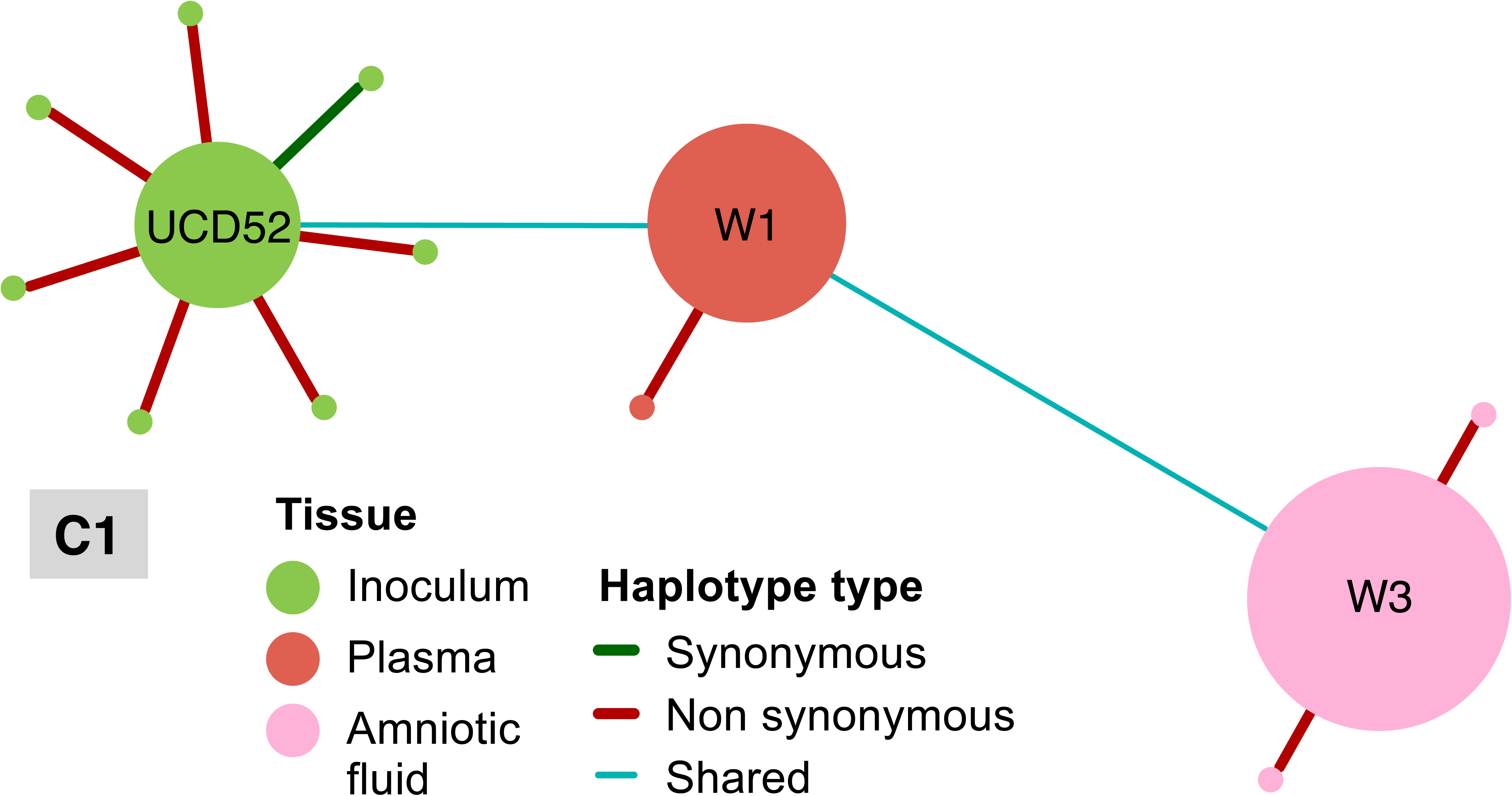
Haplotype networks for the *gB* locus across sampled tissues from the remaining 8 monkeys in the study. Colorcoding of nodes and edges are as in Figure 1, which show haplotype networks for C4, S2, and HP3. Figures S4-S7 are for monkeys C1, C2, C3, C5, respectively. Figures S8-9 are for monkeys S1 and S3 respectively. Figures S10-S11 are for monkeys HP1 and HP2, respectively.

**Figure S5.**
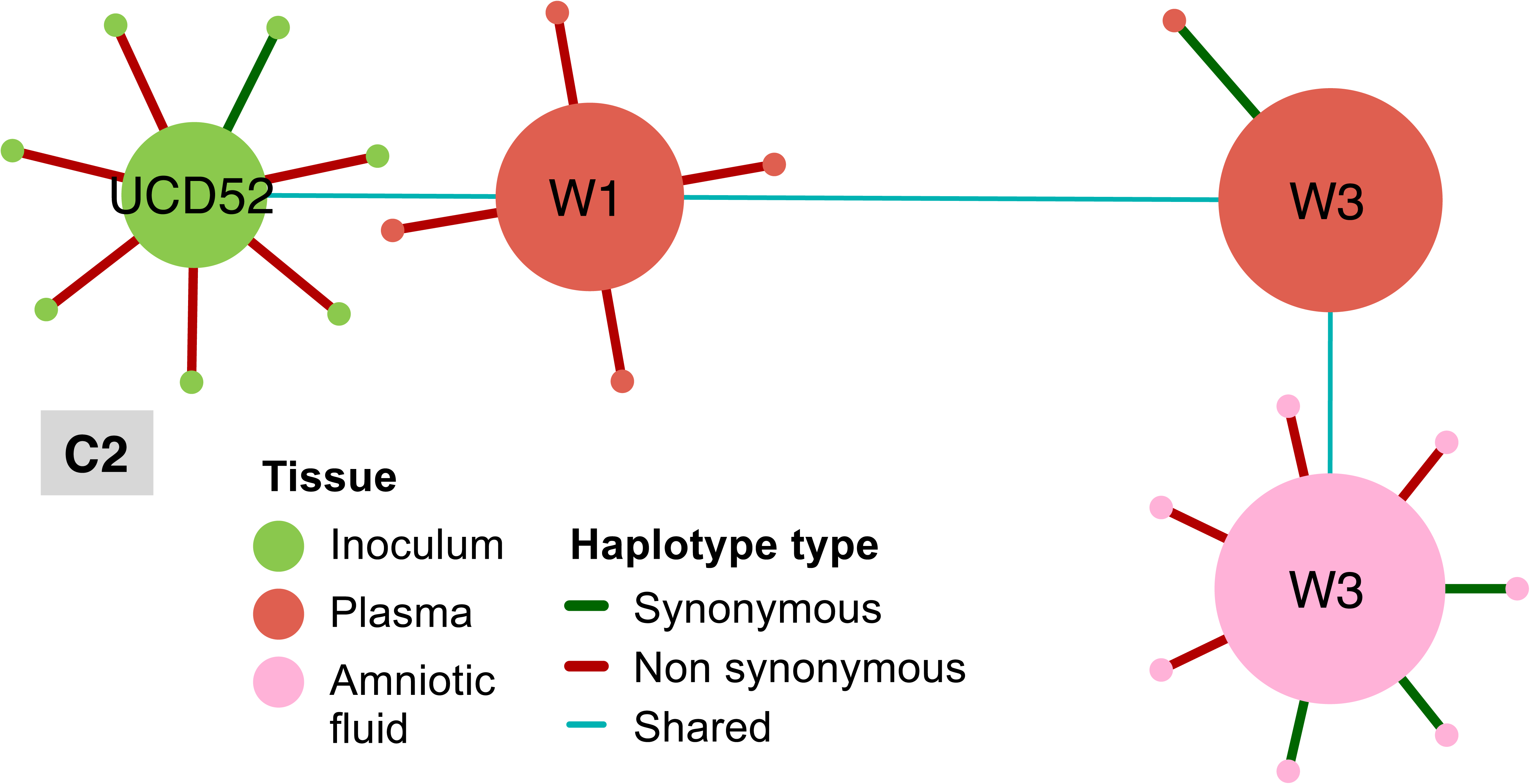
Haplotype networks for the *gB* locus across sampled tissues from the remaining 8 monkeys in the study. Colorcoding of nodes and edges are as in Figure 1, which show haplotype networks for C4, S2, and HP3. Figures S4-S7 are for monkeys C1, C2, C3, C5, respectively. Figures S8-9 are for monkeys S1 and S3 respectively. Figures S10-S11 are for monkeys HP1 and HP2, respectively.

**Figure S6.**
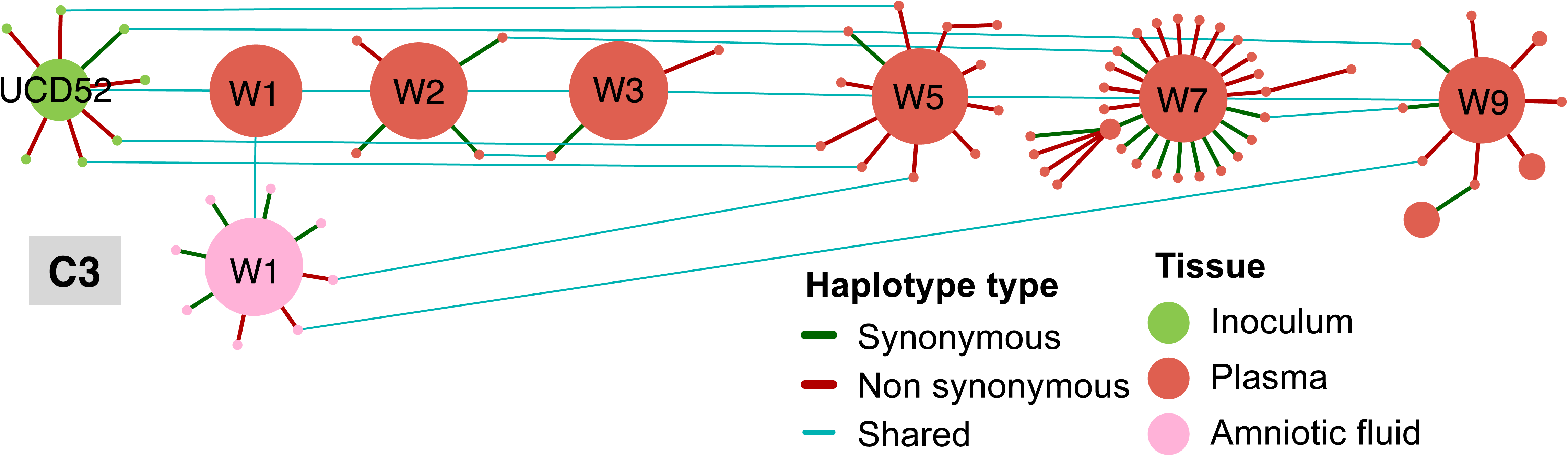
Haplotype networks for the *gB* locus across sampled tissues from the remaining 8 monkeys in the study. Colorcoding of nodes and edges are as in Figure 1, which show haplotype networks for C4, S2, and HP3. Figures S4-S7 are for monkeys C1, C2, C3, C5, respectively. Figures S8-9 are for monkeys S1 and S3 respectively. Figures S10-S11 are for monkeys HP1 and HP2, respectively.

**Figure S7.**
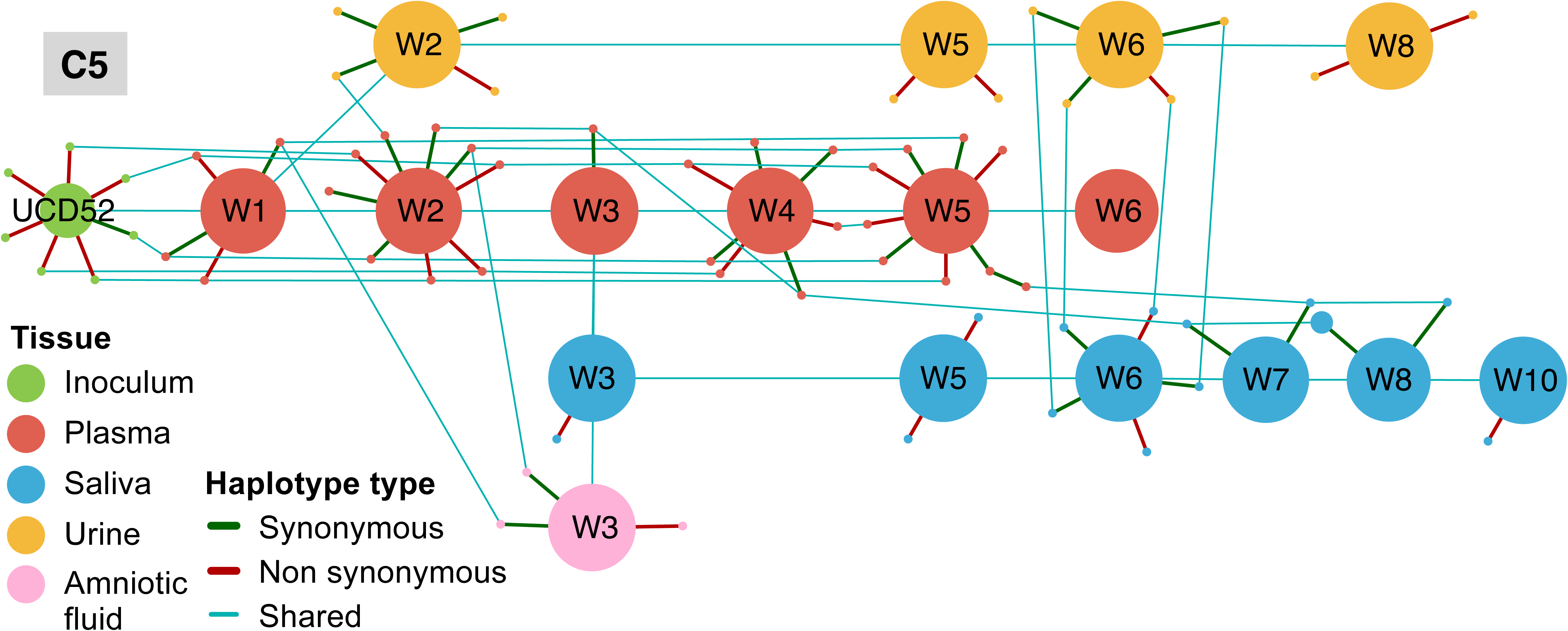
Haplotype networks for the *gB* locus across sampled tissues from the remaining 8 monkeys in the study. Colorcoding of nodes and edges are as in Figure 1, which show haplotype networks for C4, S2, and HP3. Figures S4-S7 are for monkeys C1, C2, C3, C5, respectively. Figures S8-9 are for monkeys S1 and S3 respectively. Figures S10-S11 are for monkeys HP1 and HP2, respectively.

**Figure S8.**
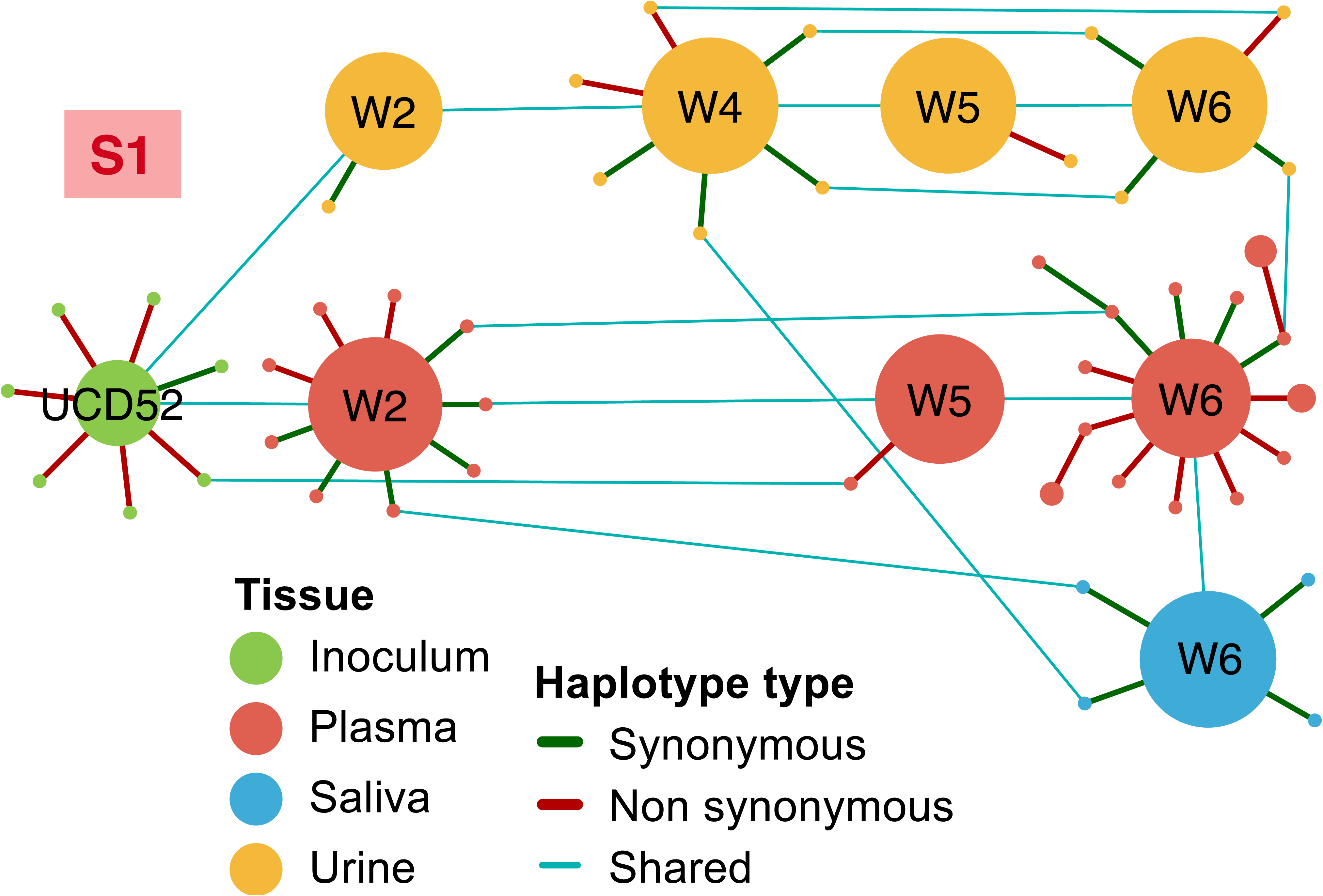
Haplotype networks for the *gB* locus across sampled tissues from the remaining 8 monkeys in the study. Colorcoding of nodes and edges are as in Figure 1, which show haplotype networks for C4, S2, and HP3. Figures S4-S7 are for monkeys C1, C2, C3, C5, respectively. Figures S8-9 are for monkeys S1 and S3 respectively. Figures S10-S11 are for monkeys HP1 and HP2, respectively.

**Figure S9.**
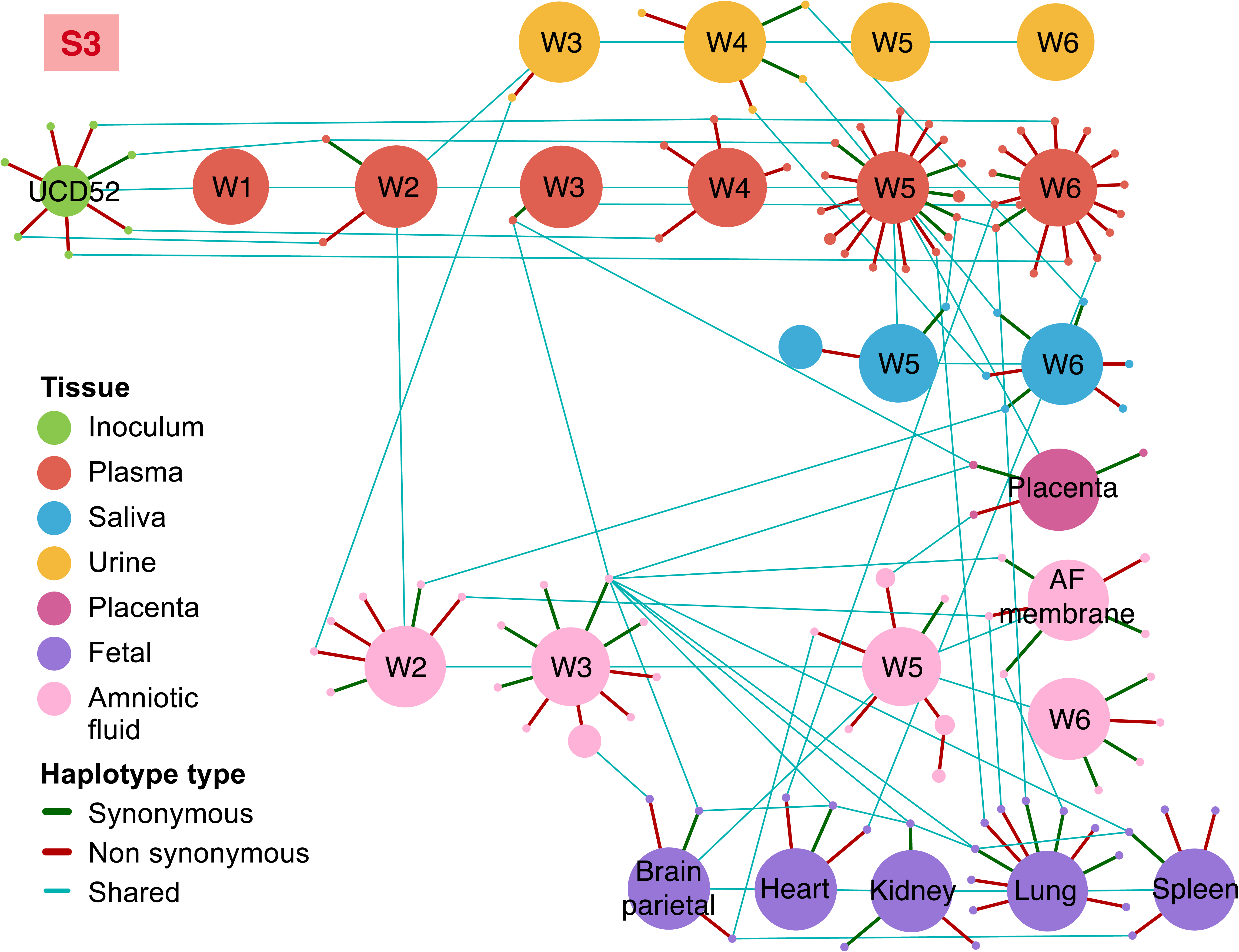
Haplotype networks for the *gB* locus across sampled tissues from the remaining 8 monkeys in the study. Colorcoding of nodes and edges are as in Figure 1, which show haplotype networks for C4, S2, and HP3. Figures S4-S7 are for monkeys C1, C2, C3, C5, respectively. Figures S8-9 are for monkeys S1 and S3 respectively. Figures S10-S11 are for monkeys HP1 and HP2, respectively.

**Figure S10.**
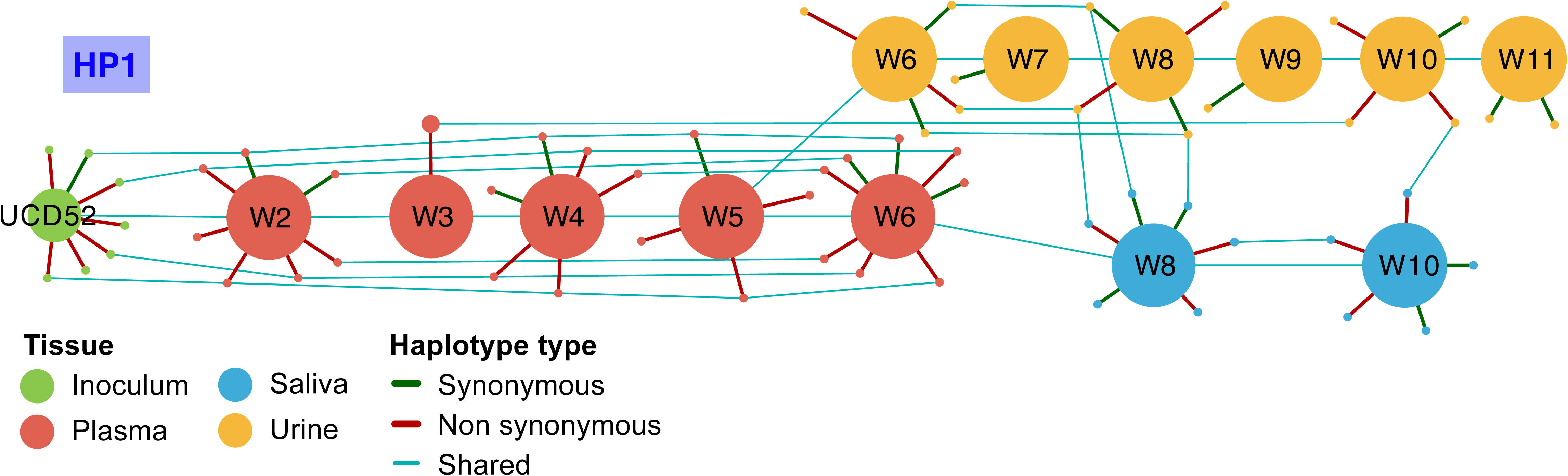
Haplotype networks for the *gB* locus across sampled tissues from the remaining 8 monkeys in the study. Colorcoding of nodes and edges are as in Figure 1, which show haplotype networks for C4, S2, and HP3. Figures S4-S7 are for monkeys C1, C2, C3, C5, respectively. Figures S8-9 are for monkeys S1 and S3 respectively. Figures S10-S11 are for monkeys HP1 and HP2, respectively.

**Figure S11.**
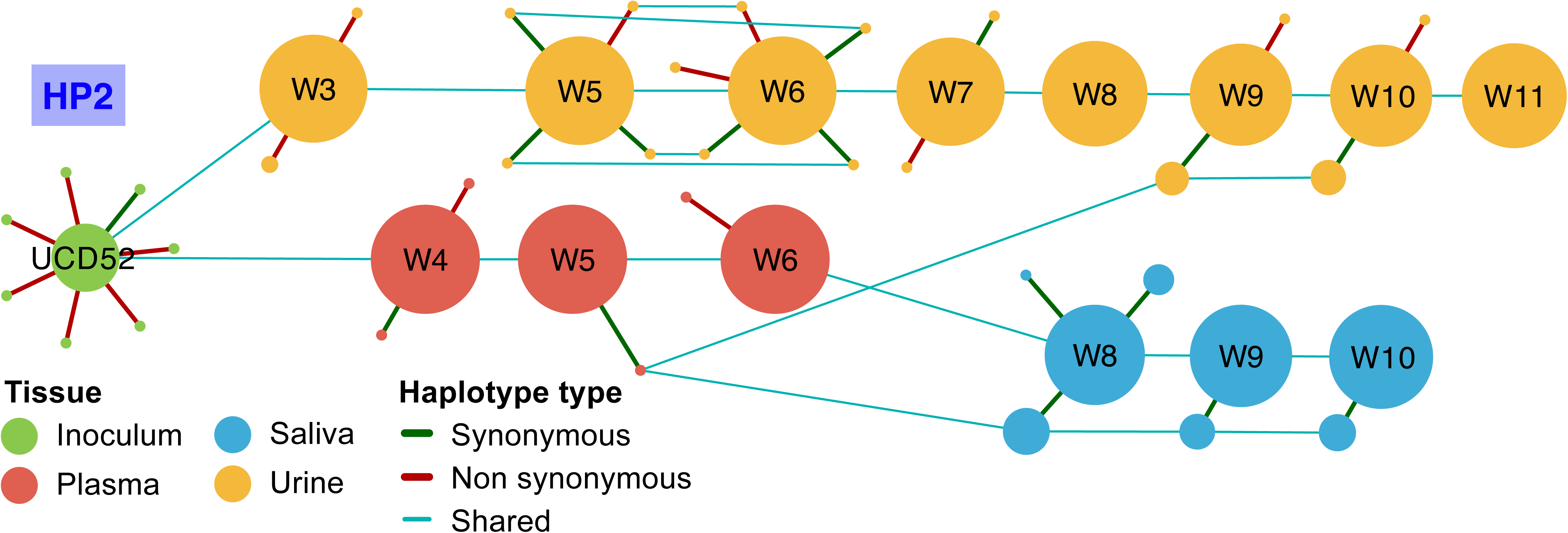
Haplotype networks for the *gB* locus across sampled tissues from the remaining 8 monkeys in the study. Colorcoding of nodes and edges are as in Figure 1, which show haplotype networks for C4, S2, and HP3. Figures S4-S7 are for monkeys C1, C2, C3, C5, respectively. Figures S8-9 are for monkeys S1 and S3 respectively. Figures S10-S11 are for monkeys HP1 and HP2, respectively.

**Figure S12.**
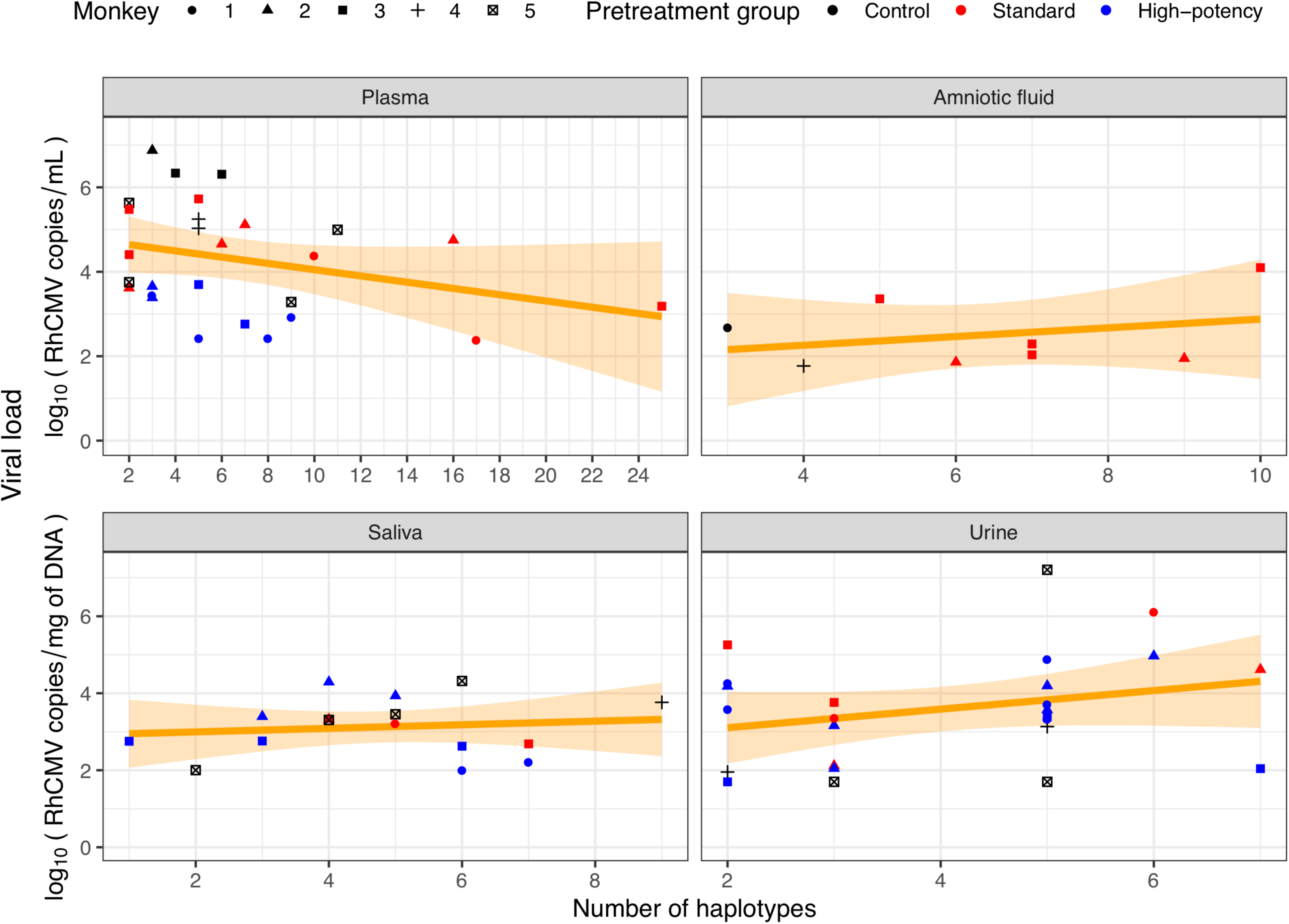
The relationship between viral load and the number of *gB* haplotypes found in each sample. The correlation between viral load and the number of *gB* haplotypes was not significantly positive for any of the four analyzed compartments (plasma, amniotic fluid, saliva, urine).

**Figure S13.**
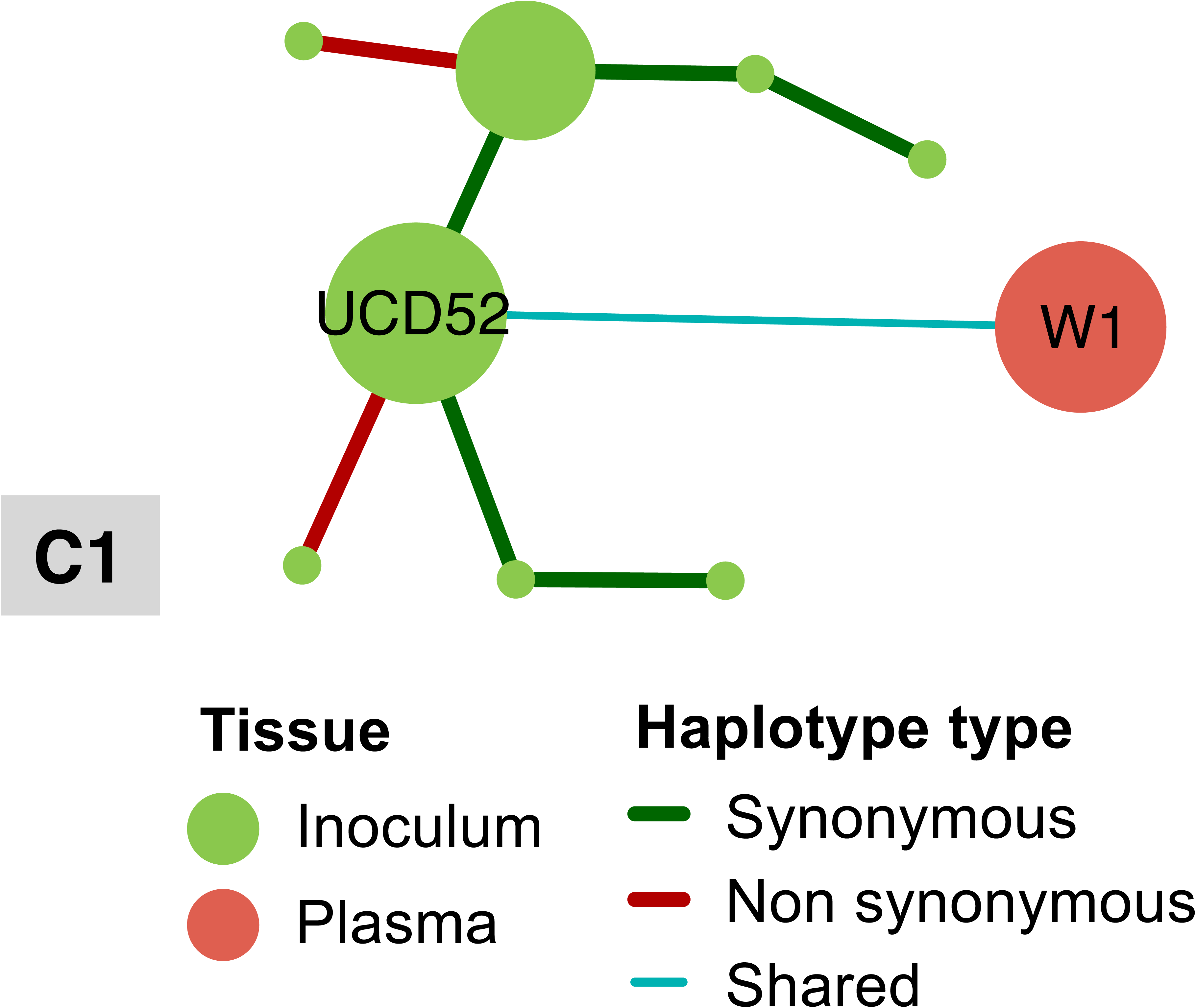
Haplotype networks for the *gL* locus across sampled tissues from each of the 10 monkeys in the study that had at least one successfully sequenced *gL* sample. Colorcoding of nodes and edges are as in Figure 1. Figures S13-S16 are for monkeys C1, C3, C4, and C5, respectively (monkey C2 did not have a successfully sequenced *gL* sample). Figures S17-S19 are for monkeys S1-S3 respectively. Figures S20-S22 are for monkeys HP1-HP3 respectively.

**Figure S14.**
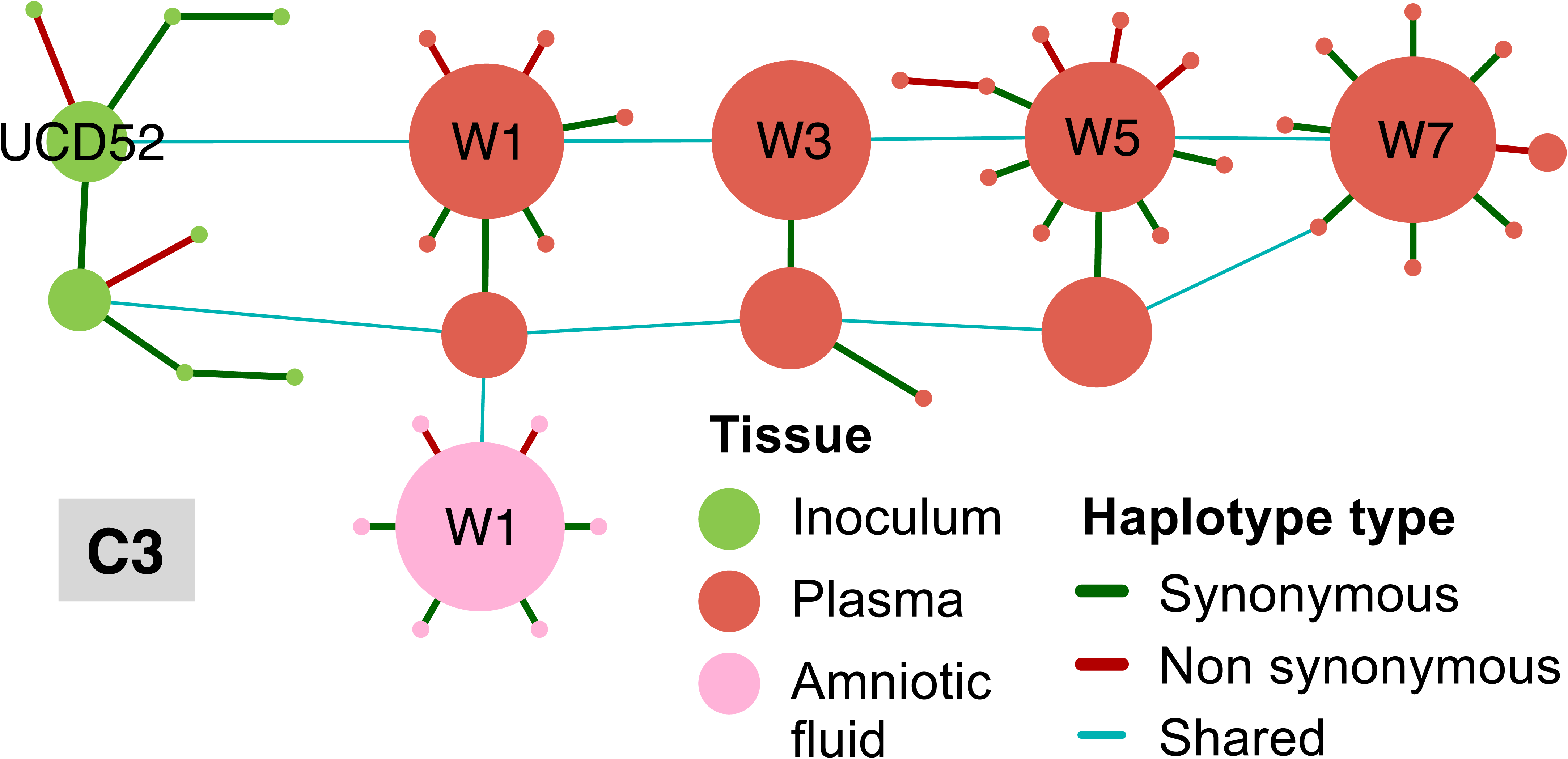
Haplotype networks for the *gL* locus across sampled tissues from each of the 10 monkeys in the study that had at least one successfully sequenced *gL* sample. Colorcoding of nodes and edges are as in Figure 1. Figures S13-S16 are for monkeys C1, C3, C4, and C5, respectively (monkey C2 did not have a successfully sequenced *gL* sample). Figures S17-S19 are for monkeys S1-S3 respectively. Figures S20-S22 are for monkeys HP1-HP3 respectively.

**Figure S15.**
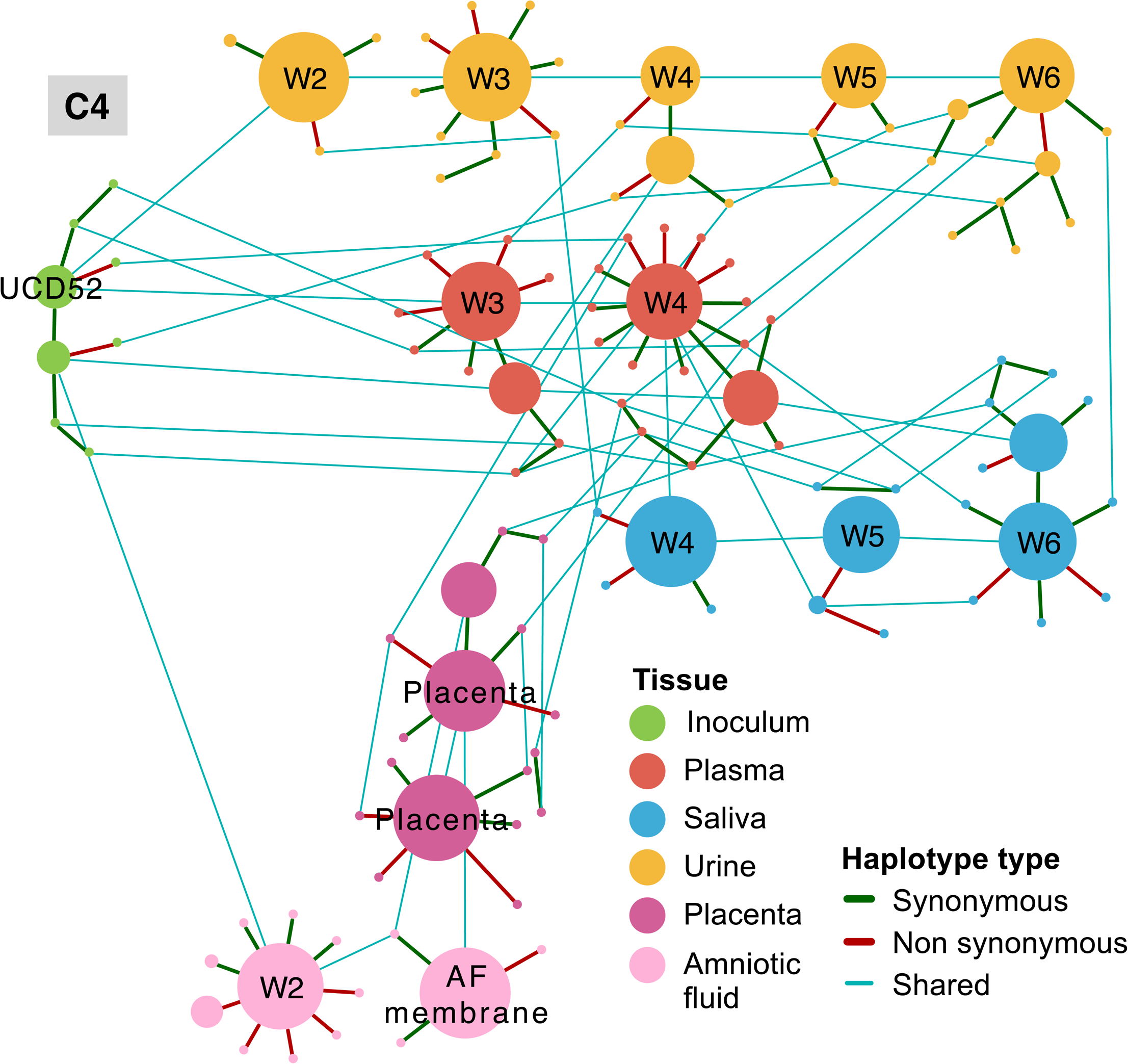
Haplotype networks for the *gL* locus across sampled tissues from each of the 10 monkeys in the study that had at least one successfully sequenced *gL* sample. Colorcoding of nodes and edges are as in Figure 1. Figures S13-S16 are for monkeys C1, C3, C4, and C5, respectively (monkey C2 did not have a successfully sequenced *gL* sample). Figures S17-S19 are for monkeys S1-S3 respectively. Figures S20-S22 are for monkeys HP1-HP3 respectively.

**Figure S16.**
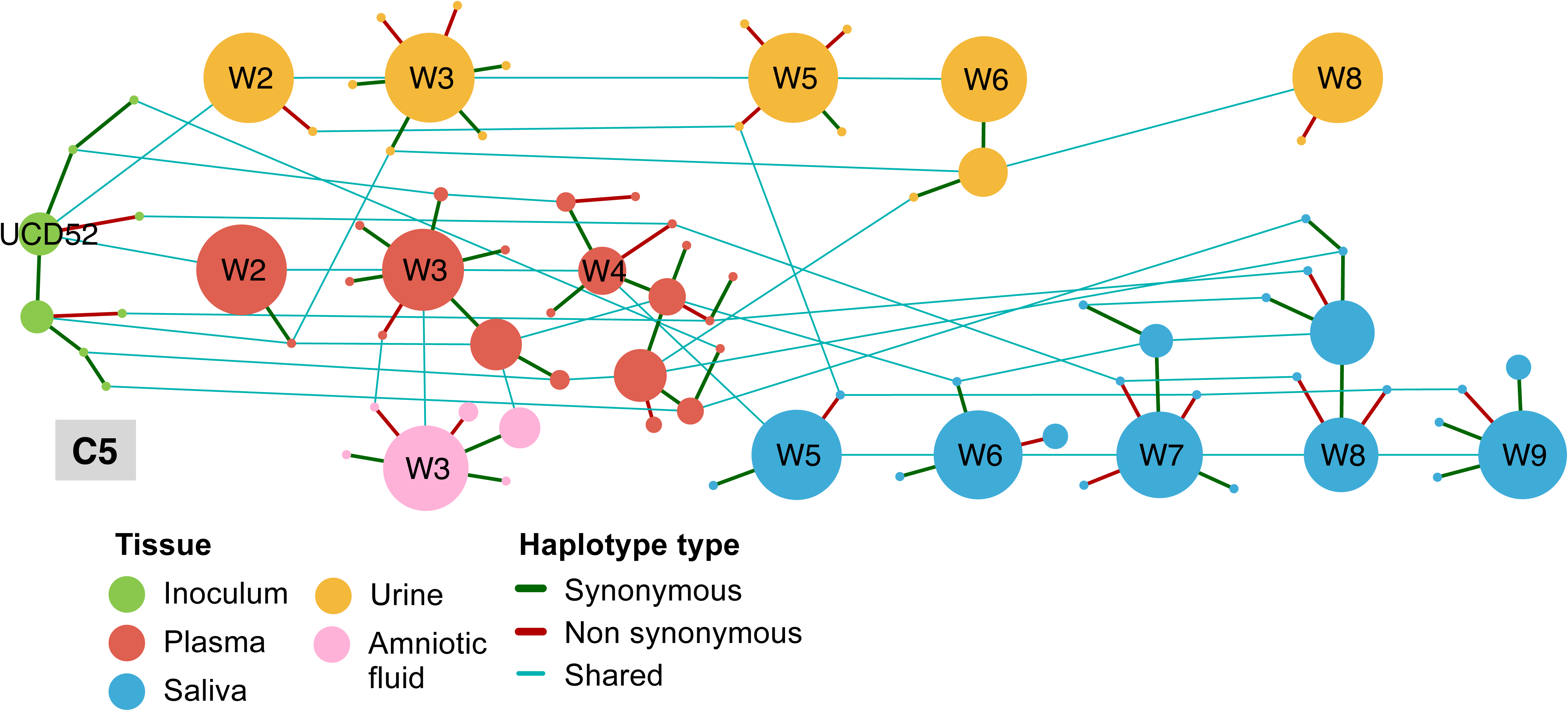
Haplotype networks for the *gL* locus across sampled tissues from each of the 10 monkeys in the study that had at least one successfully sequenced *gL* sample. Colorcoding of nodes and edges are as in Figure 1. Figures S13-S16 are for monkeys C1, C3, C4, and C5, respectively (monkey C2 did not have a successfully sequenced *gL* sample). Figures S17-S19 are for monkeys S1-S3 respectively. Figures S20-S22 are for monkeys HP1-HP3 respectively.

**Figure S17.**
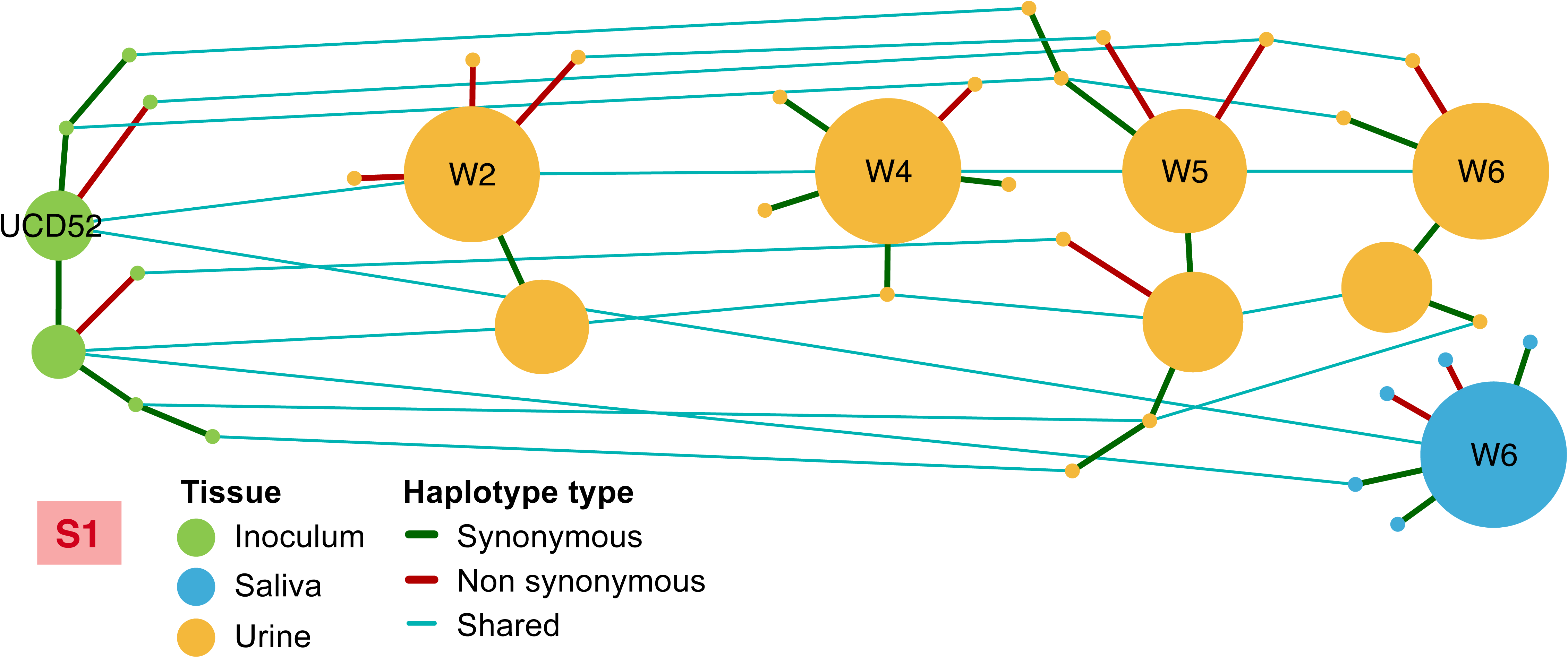
Haplotype networks for the *gL* locus across sampled tissues from each of the 10 monkeys in the study that had at least one successfully sequenced *gL* sample. Colorcoding of nodes and edges are as in Figure 1. Figures S13-S16 are for monkeys C1, C3, C4, and C5, respectively (monkey C2 did not have a successfully sequenced *gL* sample). Figures S17-S19 are for monkeys S1-S3 respectively. Figures S20-S22 are for monkeys HP1-HP3 respectively.

**Figure S18.**
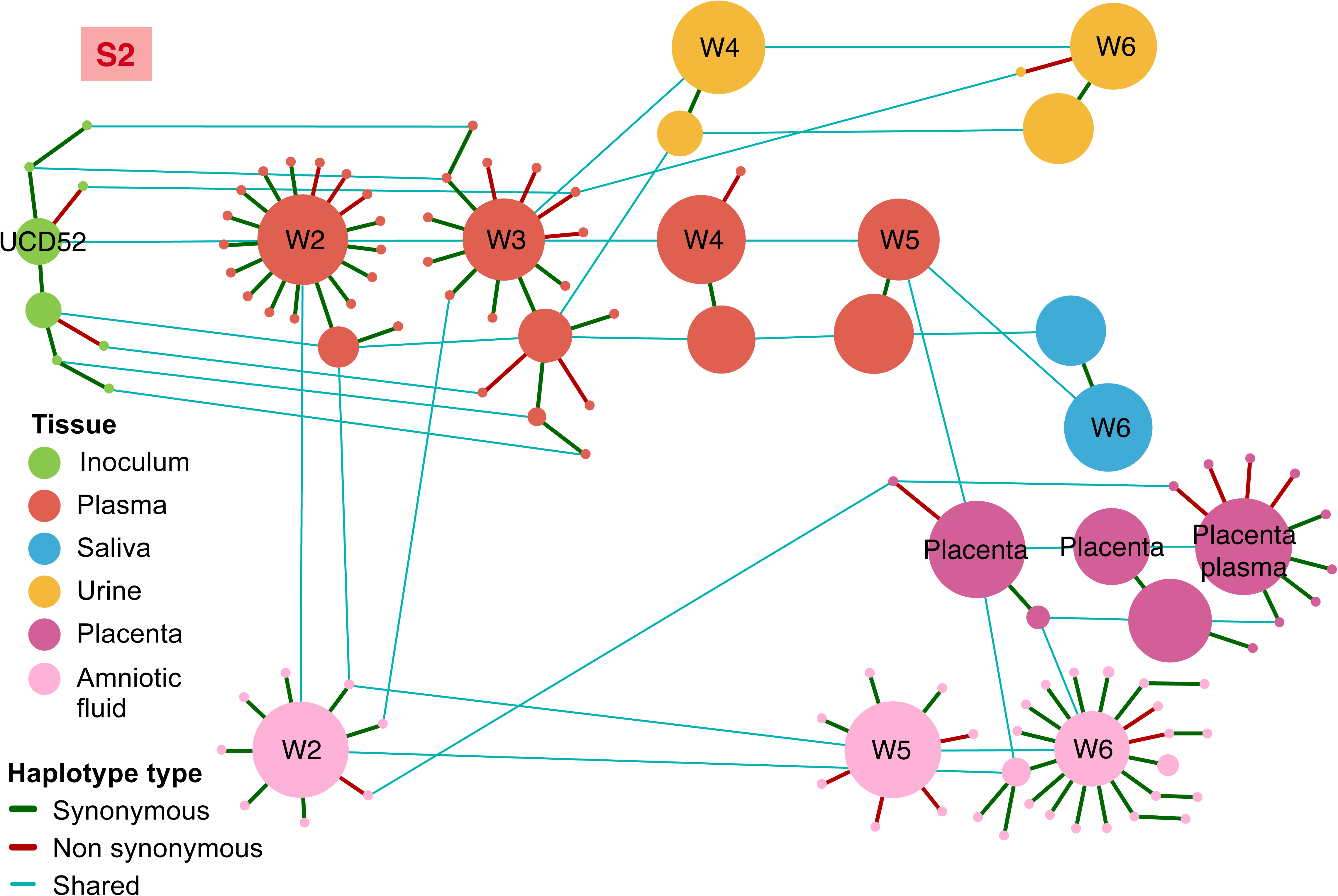
Haplotype networks for the *gL* locus across sampled tissues from each of the 10 monkeys in the study that had at least one successfully sequenced *gL* sample. Colorcoding of nodes and edges are as in Figure 1. Figures S13-S16 are for monkeys C1, C3, C4, and C5, respectively (monkey C2 did not have a successfully sequenced *gL* sample). Figures S17-S19 are for monkeys S1-S3 respectively. Figures S20-S22 are for monkeys HP1-HP3 respectively.

**Figure S19.**
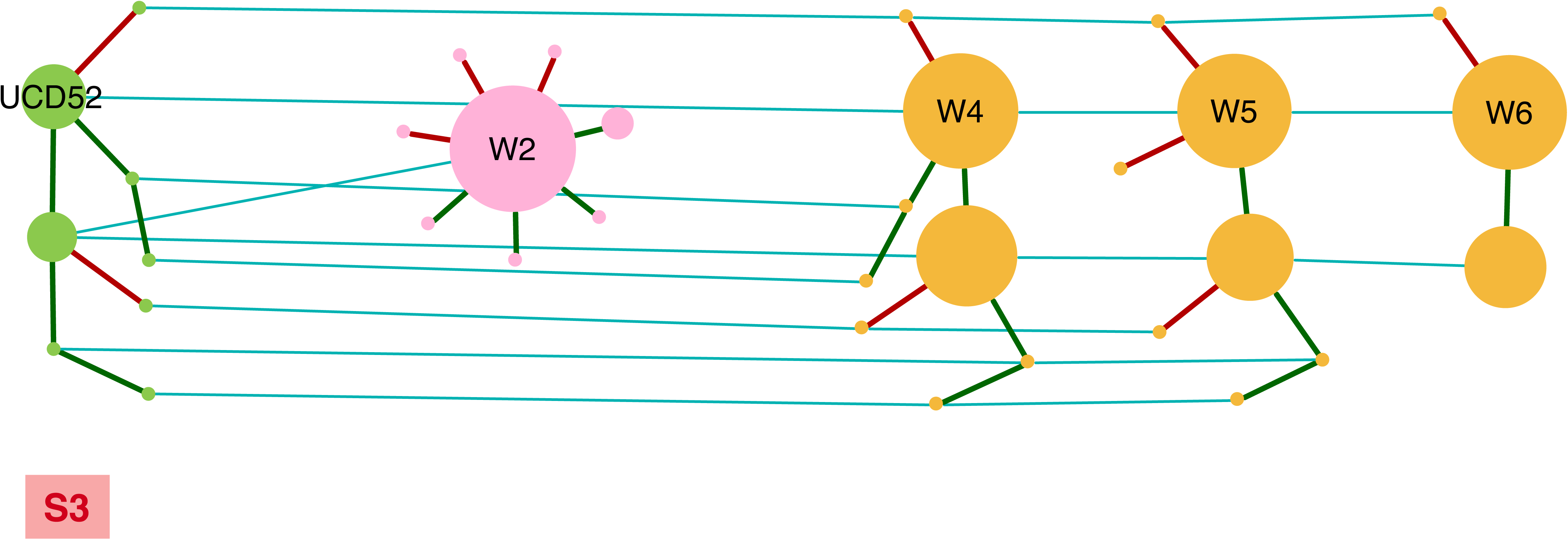
Haplotype networks for the *gL* locus across sampled tissues from each of the 10 monkeys in the study that had at least one successfully sequenced *gL* sample. Colorcoding of nodes and edges are as in Figure 1. Figures S13-S16 are for monkeys C1, C3, C4, and C5, respectively (monkey C2 did not have a successfully sequenced *gL* sample). Figures S17-S19 are for monkeys S1-S3 respectively. Figures S20-S22 are for monkeys HP1-HP3 respectively.

**Figure S20.**
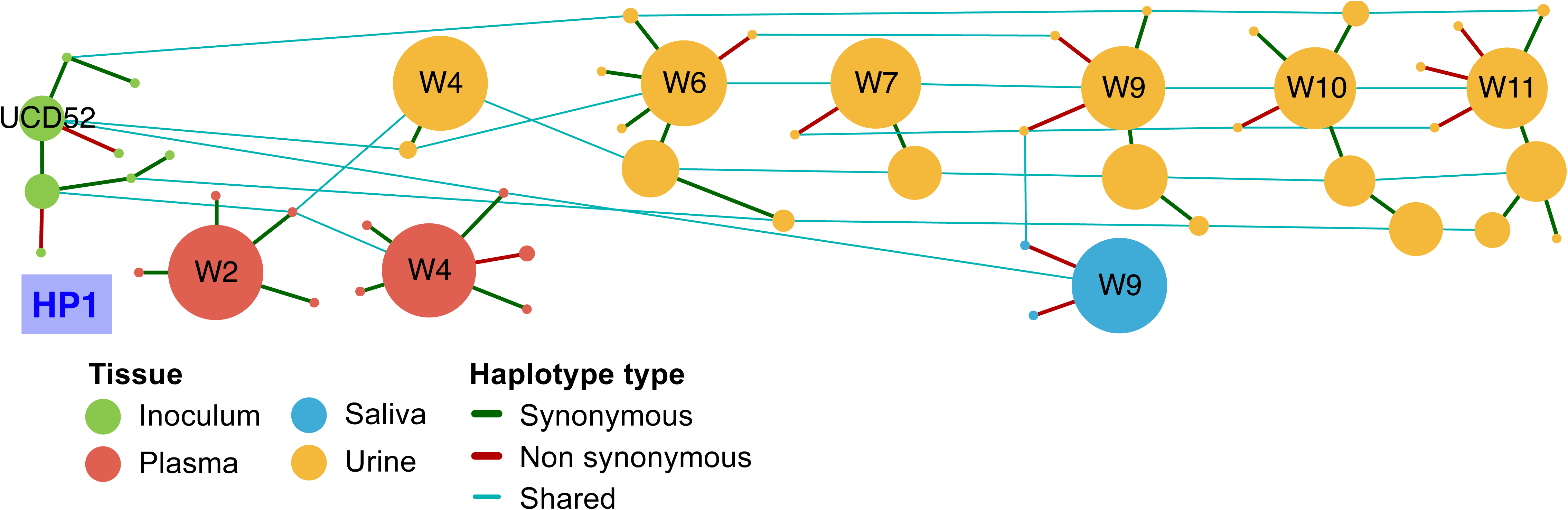
Haplotype networks for the *gL* locus across sampled tissues from each of the 10 monkeys in the study that had at least one successfully sequenced *gL* sample. Colorcoding of nodes and edges are as in Figure 1. Figures S13-S16 are for monkeys C1, C3, C4, and C5, respectively (monkey C2 did not have a successfully sequenced *gL* sample). Figures S17-S19 are for monkeys S1-S3 respectively. Figures S20-S22 are for monkeys HP1-HP3 respectively.

**Figure S21.**
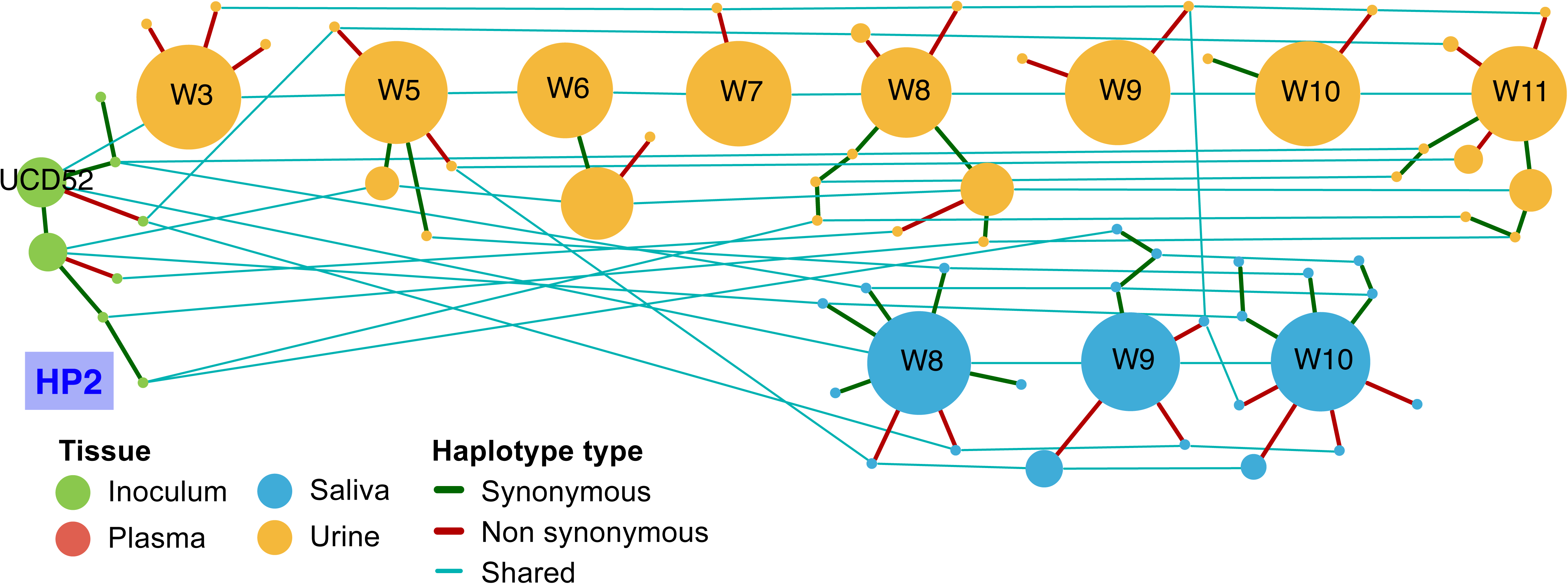
Haplotype networks for the *gB* locus across sampled tissues from each of the 10 monkeys in the study that had at least one successfully sequenced *gL* sample. Colorcoding of nodes and edges are as in Figure 1. Figures S13-S16 are for monkeys C1, C3, C4, and C5, respectively (monkey C2 did not have a successfully sequenced *gL* sample). Figures S17-S19 are for monkeys S1-S3 respectively. Figures S20-S22 are for monkeys HP1-HP3 respectively.

**Figure S22.**
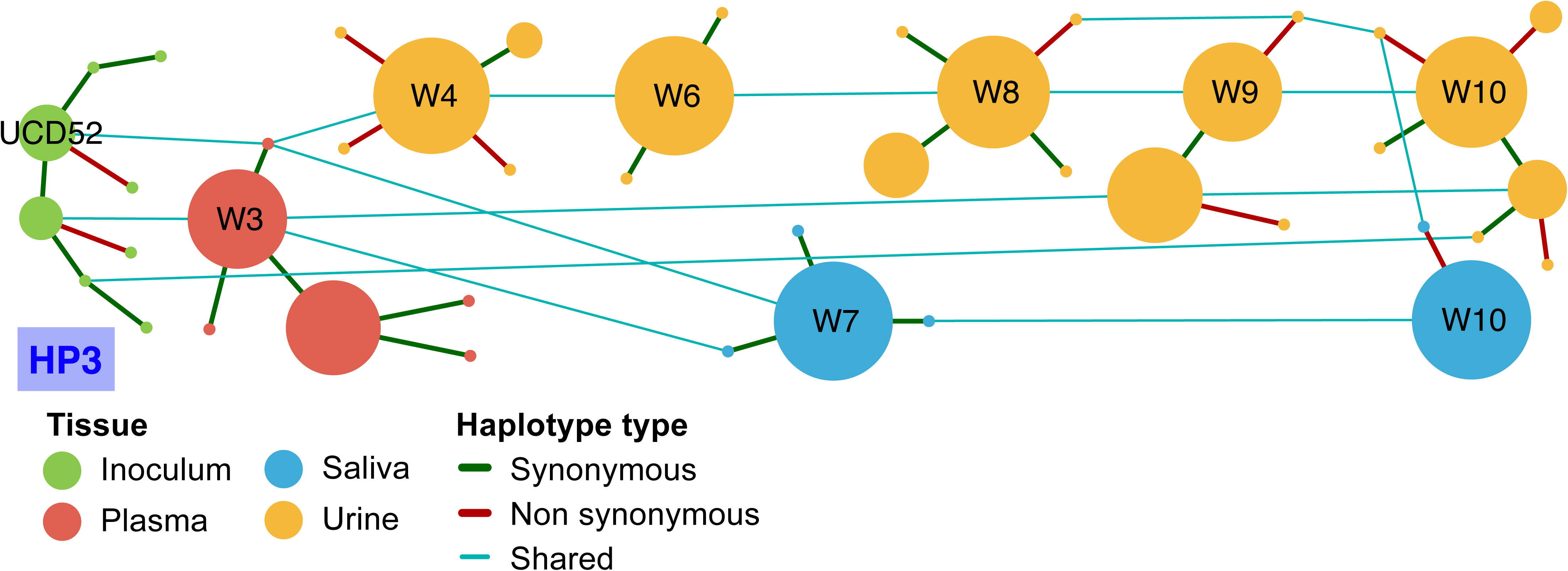
Haplotype networks for the *gL* locus across sampled tissues from each of the 10 monkeys in the study that had at least one successfully sequenced *gL* sample. Colorcoding of nodes and edges are as in Figure 1. Figures S13-S16 are for monkeys C1, C3, C4, and C5, respectively (monkey C2 did not have a successfully sequenced *gL* sample). Figures S17-S19 are for monkeys S1-S3 respectively. Figures S20-S22 are for monkeys HP1-HP3 respectively.

**Figure S23.**
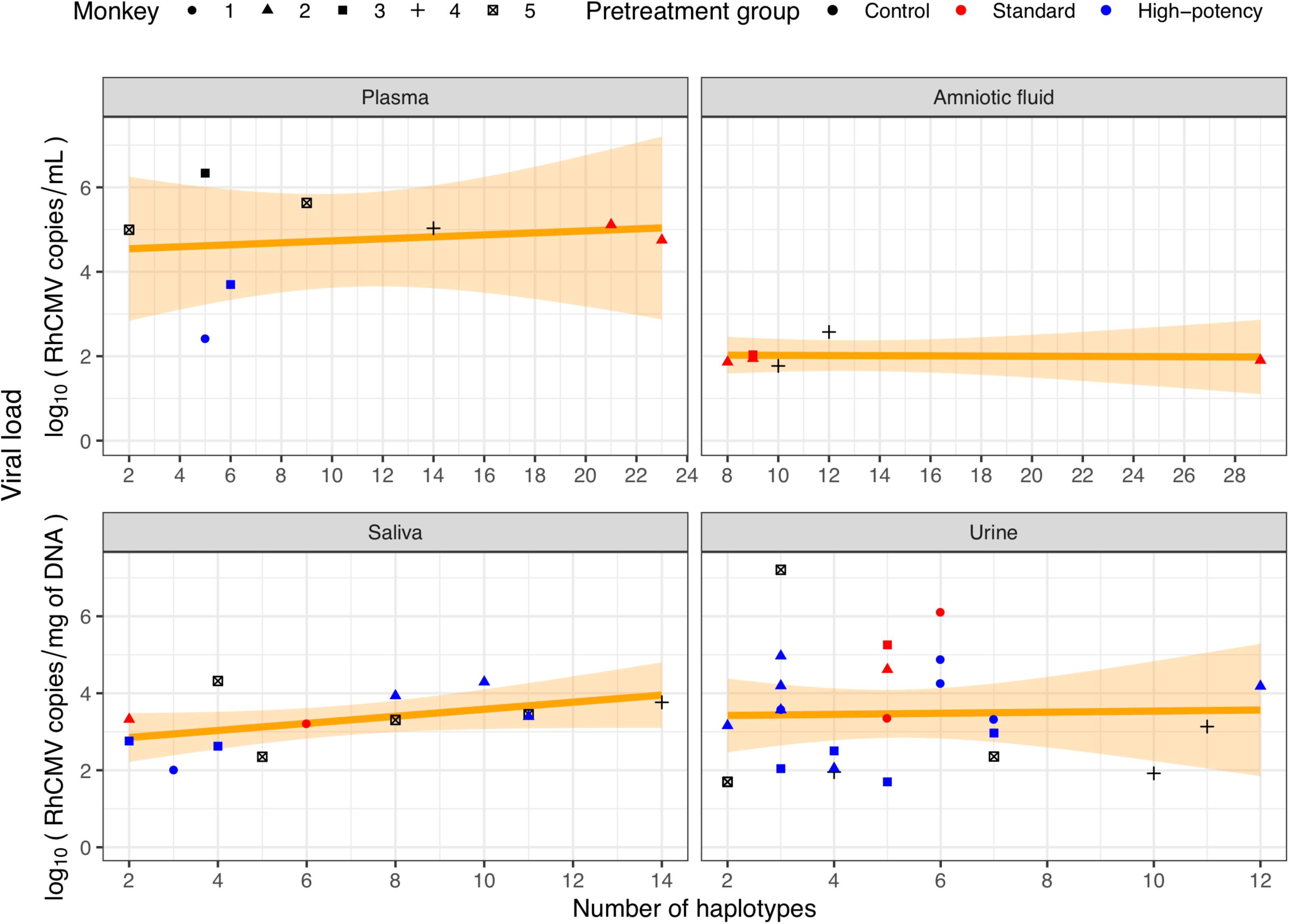
The relationship between viral load and the number of *gL* haplotypes found in each samples. The correlation between viral load and the number of *gL* haplotypes was not significantly positive for any of the four analyzed compartments (plasma, amniotic fluid, saliva, urine).

**Figure S24.**
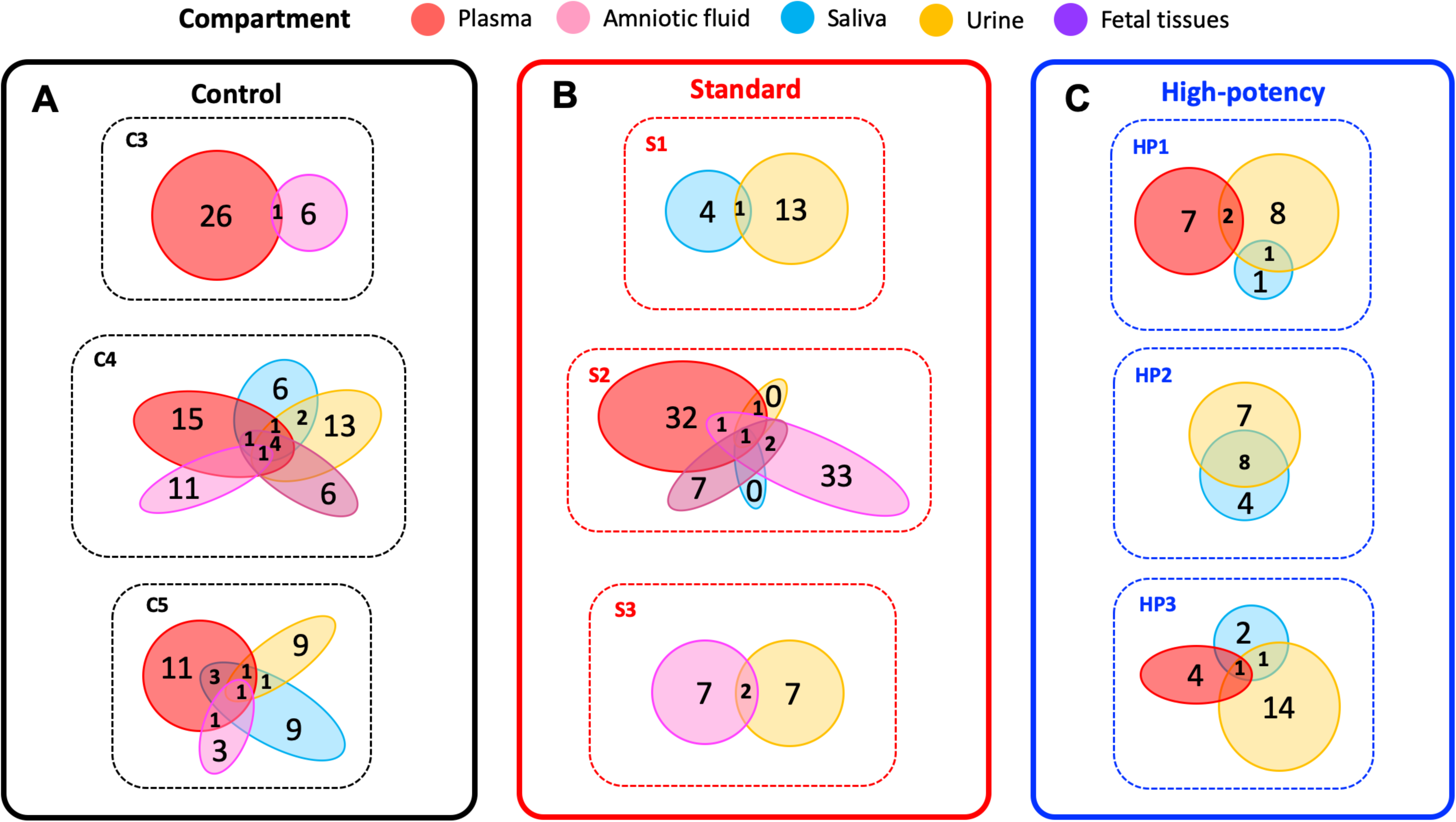
The number of UCD52 minor *gL* haplotypes that are either shared or unique across compartments, by monkey. Here, the set of minor haplotypes for a given compartment includes all timepoint samples from that compartment. Patterns of minor haplotype sharing for (A) control group monkeys, (B) standard pretreatment group monkeys, and (C) high-potency pretreatment group monkeys. Compartments are colorcoded as in Figure 1.

**Figure S25.**
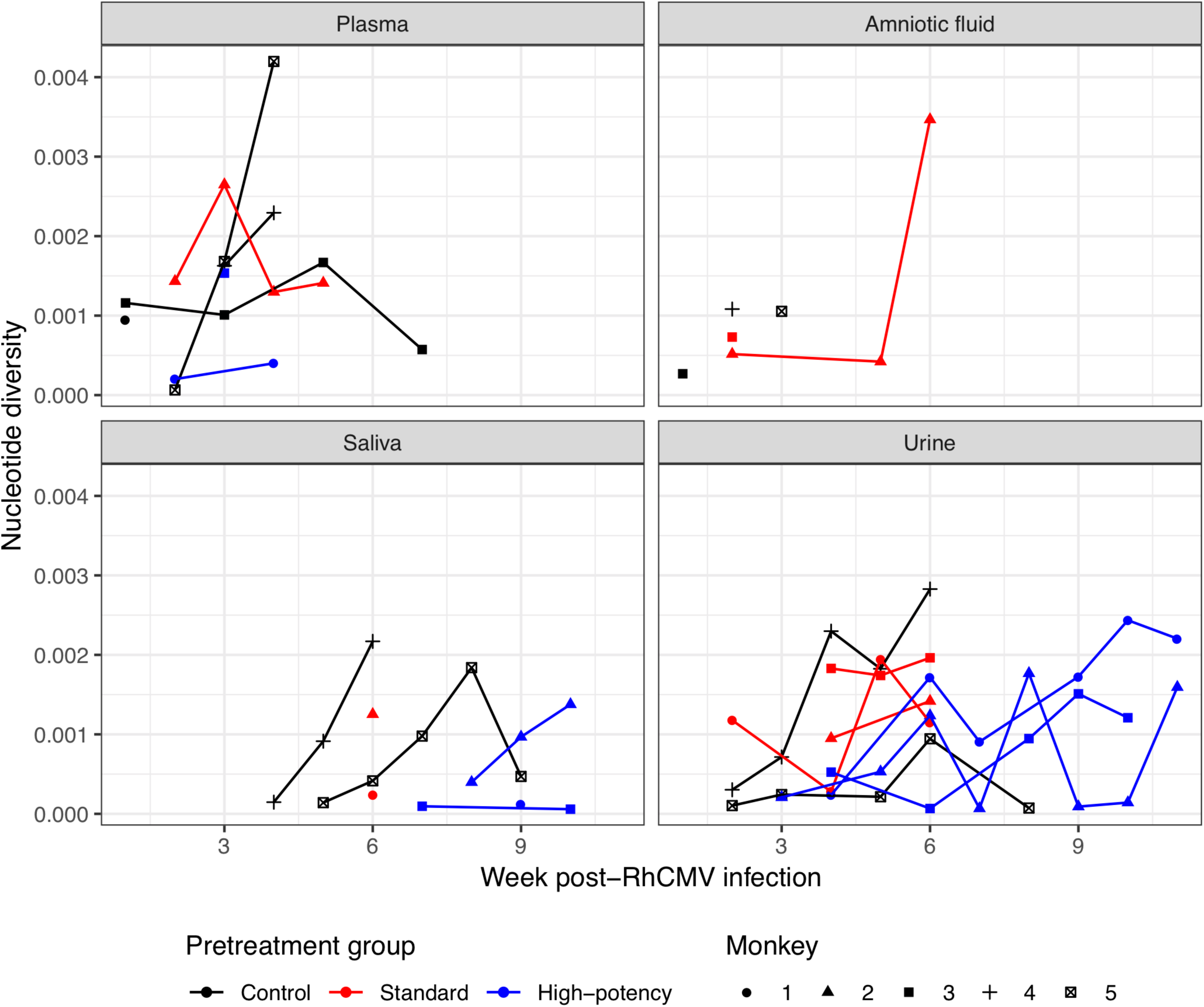
Pairwise genetic diversity *π* over time, by tissue, for the *gL* locus. Marker symbols correspond with those in Figure S2.

**Table S1.**
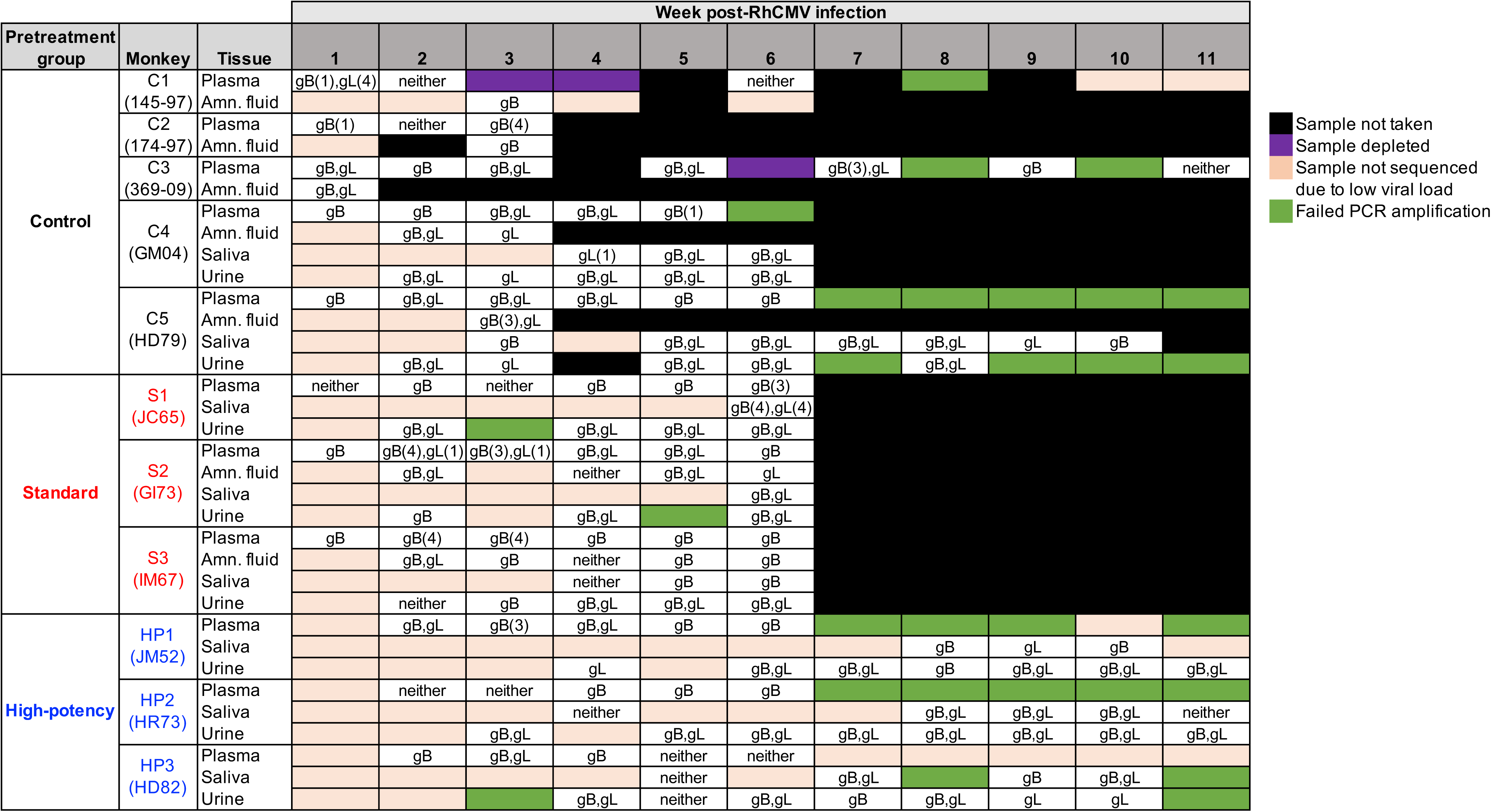
Sampling times and tissues across the 11 studied dams. Dams are separated by pretreatment group: control (C1-C5), standard (S1-S3), and high-potency (HP1-HP3). In addition to the C1-C5, S1-S3, and HP1-HP3 identifiers, individual monkeys are identified according to names previously used in [9] and [23]. Cells are colored according to the legend provided. Text in the white-colored cells indicate which loci were successfully sequenced and included in our analyses (*gB* = glycoprotein B region; *gL* = glycoprotein L region). Numbers in the cells, when present, indicate the number of sample replicates that were available for analysis, when not two.

**Table S2.** Fetal-maternal interface and fetal tissues analyzed in the study. All listed samples had two successfully sequenced replicates.

| Pretreatment group | Monkey | Tissue | Week | Samples |
| --- | --- | --- | --- | --- |
| Control | C4 | Amn. fluid membrane | 3 | gB, gL |
|  |  | Placenta -1 | 3 | gB, gL |
|  |  | Placenta -2 | 3 | gB, gL |
| Standard | S2 | Placenta - 1 | 6 | gB, gL |
|  |  | Placenta - 2 | 6 | gB, gL |
|  |  | Placenta plasma | 6 | gB, gL |
|  | S3 | Amn. fluid membrane | 6 | gB |
|  |  | Placenta - 2 | 6 | gB |
|  |  | Lung | 6 | gB |
|  |  | Brain - parietal | 6 | gB |
|  |  | Spleen | 6 | gB |
|  |  | Heart | 6 | gB |
|  |  | Kidney | 6 | gB |

## Supplementary appendices

**Appendix S1. Haplotypes information at *gB* locus.** List of all unique haplotypes identified in all the monkeys and viral stocks at the *gB* locus, including the number and type of mutations as compared to their respective strain reference.

**Appendix S2. Haplotypes information at *gL* locus.** List of all unique haplotypes identified in all the monkeys and viral stocks at the *gL* locus, including the number and type of mutations as compared to their respective strain reference.

**Appendix S3. Haplotypes per sample at *gB* locus.** Table containing all the haplotypes found in each sample at the *gB* locus. Samples are defined by the monkey ID, tissue of origin and collection week post-RhCMV infection. Each haplotype entry includes its reference strain and relative frequency in a given sample.

**Appendix S4. Haplotypes per sample at *gL* locus.** Table containing all the haplotypes found in each sample at the *gL* locus. Samples are defined by the monkey ID, tissue of origin and collection week post-RhCMV infection. Each haplotype entry includes its reference strain and relative frequency in a given sample.

## Notes

#### Summary of Updates

High-quality figures. Adding supplementary figures and tables.

